# CNN MouseNet: A biologically constrained convolutional neural network model for mouse visual cortex

**DOI:** 10.1101/2021.10.04.463077

**Authors:** Jianghong Shi, Bryan Tripp, Eric Shea-Brown, Stefan Mihalas, Michael Buice

**Affiliations:** Applied Mathematics and Computational Neuroscience Center, University of Washington, Seattle, WA, United States; Centre for Theoretical Neuroscience, University of Waterloo, Waterloo, Ontario, Canada; Allen Institute, Seattle, WA, United States

## Abstract

Convolutional neural networks trained on object recognition derive inspiration from the neural architecture of the visual system in primates, and have been used as models of the feedforward computation performed in the primate ventral stream. In contrast to the deep hierarchical organization of primates, the visual system of the mouse has a shallower arrangement. Since mice and primates are both capable of visually guided behavior, this raises questions about the role of architecture in neural computation. In this work, we introduce a novel framework for building a biologically constrained convolutional neural network model of the mouse visual cortex. The architecture and structural parameters of the network are derived from experimental measurements, specifically the 100-micrometer resolution interareal connectome, the estimates of numbers of neurons in each area and cortical layer, and the statistics of connections between cortical layers. This network is constructed to support detailed task-optimized models of mouse visual cortex, with neural populations that can be compared to specific corresponding populations in the mouse brain. Using a well-studied image classification task as our working example, we demonstrate the computational capability of this mouse-sized network. Given its relatively small size, MouseNet achieves roughly 2/3rds the performance level on ImageNet as VGG16. In combination with the large scale Allen Brain Observatory Visual Coding dataset, we use representational similarity analysis to quantify the extent to which MouseNet recapitulates the neural representation in mouse visual cortex. Importantly, we provide evidence that optimizing for task performance does not improve similarity to the corresponding biological system beyond a certain point. We demonstrate that the distributions of some physiological quantities are closer to the observed distributions in the mouse brain after task training. We encourage the use of the MouseNet architecture by making the code freely available.

**Author summary:** Task-driven deep neural networks have shown great potential in predicting functional responses of biological neurons. Nevertheless, they are not precise biological analogues, raising questions about how they should be interpreted. Here, we build new deep neural network models of the mouse visual cortex (MouseNet) that are biologically constrained in detail, not only in terms of the basic structure of their connectivity, but also in terms of the count and hence density of neurons within each area, and the spatial extent of their projections. Equipped with the MouseNet model, we can address key questions about mesoscale brain architecture and its role in task learning and performance.We ask, and provide a first set of answers, to: What is the performance of a mouse brain-sized – and mouse brain-structured – model on benchmark image classification tasks? How does the training of a network on this task affect the functional properties of specified layers within the biologically constrained architecture – both overall, and in comparison with recorded function of mouse neurons? We anticipate much future work on allied questions, and the development of more sophisticated models in both mouse and other species, based on the freely available MouseNet model and code which we develop and provide here.

## Introduction

Convolutional neural networks (CNNs) trained on object recognition derive some inspiration from the neuroscience of the visual system in primates, and have been used as models of feedforward computation performed in the primate ventral stream [1–3]. Indeed, the activity in these networks often resembles activity recorded from areas of the primate visual system, from oriented Gabor-like features in early layers [4] to responses to curves and more complex geometries [5] and even functional, or *representational,* similarity at the population level [6,7]. Task-trained artificial neural networks have been shown to produce similar neural representations or develop predictive models of neural activity in visual [8–10], auditory [11], rodent whisker areas [12], and more [13–15]. Despite these successes and the clear power of CNNs to solve machine learning problems in the visual domain, among others [4,16], they are not structural or architectural analogues for the underlying biological circuits. Recent endeavors [17,18] show that imposing brain like structure such as shallowness and recurrence in network models can improve their functional similarity to the primate brain. The interplay of architecture and computation remains an open problem in both machine learning and neuroscience.

This issue is especially pronounced for studies of mouse visual cortex, a field undergoing explosive growth. Large scale tract tracing data sets have revealed neuro-anatomical structure in unprecedented detail [19–22]. From these efforts we have learned, in contrast to the hierarchical organization of primates, that the visual system of the mouse has a much more parallel structure [23]. Since rodents are capable of visually guided behavior including invariant object recognition [24, 25], this raises questions about the role of architecture in neural computation. Recently, data from a large-scale physiological survey of neural activity in the mouse visual system [26] was used to compare the representations of visual stimuli in cortex with those of modern deep networks [27–29]. It was found that even purportedly “early” regions such as V1 in mouse cortex are more similar at the level of representation to middle layers of networks such as VGG16, rather than to early layers that respond optimally to simple visual features and bear more resemblance to the “simple” and “complex” cells normally supposed to describe the early visual pathway. However, the profound difference in architecture between modern CNNs and the mouse cortex raises significant challenges in interpreting these findings. To begin, many modern computational models of vision (in particular CNNs, which often have a high input resolution) have a larger number of units than the number of neurons in mouse visual cortex. Moreover, CNNs from computational vision are largely of feedforward type, either purely so or with some skip connections (e.g., in ResNet architectures), which ignores the large amount of recurrence present in real neural circuits. Furthermore, the mouse thalamo-cortical system is quite shallow [23]. Most importantly, as stated above and detailed more below, the mouse visual cortex has an intriguing parallel structure.

Here we introduce a novel framework for incorporating these data to build a biologically constrained convolutional neural network model of the mouse visual cortex — the CNN MouseNet. Convolutional neural networks share weights across the visual field, and thus form an approximation of the functional properties that may be imposed by translation invariance of natural stimuli leading to equivariant representations in neural systems [1–3]. This weight sharing makes them much easier to train, which is an important practical consideration for model development. The structural parameters of MouseNet are derived from experimental measurements, specifically estimates of numbers of neurons in each area and cortical layer, the 100-micrometer resolution interareal connectome, and the statistics of connections between cortical layers.

MouseNet is constructed to support detailed task-optimized models of mouse visual cortex, with neural populations that can be compared to specific corresponding populations in the mouse brain. To demonstrate the usage of MouseNet, we use standard image classification tasks as working examples; specifically, we train MouseNet to perform classification using the ImageNet Large Scale Visual Recognition Challenge 2012 (ILSVRC2012) [30] as well as the CIFAR10 [31] data sets.

We find that, although MouseNet is much smaller than a typical CNN and has specific architectural differences, it can reach above 90% validation accuracy on CIFAR10, and roughly 2/3rds of the performance level of a typical CNN (VGG16) on the more challenging ImageNet classification benchmark.

Next, using the large-scale functional data sets from the Allen Brain Observatory [26] on visual responses of neurons across visual cortex, we investigate the functional properties of the MouseNet architecture after training on the ImageNet dataset. We use representational similarity analysis [27, 32, 33] to investigate the relative effects of task-training on the different computational layers in the model. We see that ImageNet classification training of MouseNet makes responses in its corresponding level of layers more similar to responses recorded from the mouse brain.

We then contrast these results for the biologically constrained MouseNet with those for a standard CNN network, VGG16, trained on the same task. We show that the representational similarity of MouseNet to the mouse brain is comparable to that of VGG16, even though VGG16 produces significantly higher task performance.

We study the training process for both networks, and find that the highest SSM values between a model neural network and the mouse brain areas are not necessarily achieved by the best performing models, rather at early or intermediate points during the training process. We take this as an indication that image classification using ImageNet is not the appropriate task to describe the mouse visual cortex (or at least those regions measured in the Allen Brain Observatory) rather than a failure of the task-training approach. This conclusion is perhaps to be expected. However, we feel that the use of object recognition is an important baseline in comparison with established results in primate.

Furthermore, in addition to broad measures of representational similarity across images, we also demonstrate the effect of image classification training on MouseNet by showing how it affects the other functional properties such as lifetime sparseness and orientation selectivity index [26]. We find that training drives both of these properties to become more similar between MouseNet and the biological mouse brain. Finally, by comparing both VGG16 and MouseNet representations in individual layers before and after training, we find that the image classification task makes MouseNet layers more diverse after training, a phenomenon we attribute to the parallel pathways in the MouseNet architecture.

Overall, we describe an open framework for constructing MouseNet that is general and can be easily modified to incorporate new data on the structure of the mouse brain [34]. Likewise, MouseNet can be readily trained on other tasks, including those corresponding more closely to natural behavior. We encourage future research along these lines by making the Python code publicly available at https://github.com/mabuice/Mouse_CNN, together with the step-by-step description of the model construction that we present next.

## Construction of CNN MouseNet

In this section, we introduce our framework for constructing the CNN MouseNet. Fig 1 shows an overview of this framework. The basic idea is to use available sources of anatomical data (e.g. tract tracing data, cell counts, and statistics of short-range connections) to constrain the CNN network structure and architectural hyperparameters. We discuss the details of each step below.

**Fig 1.**
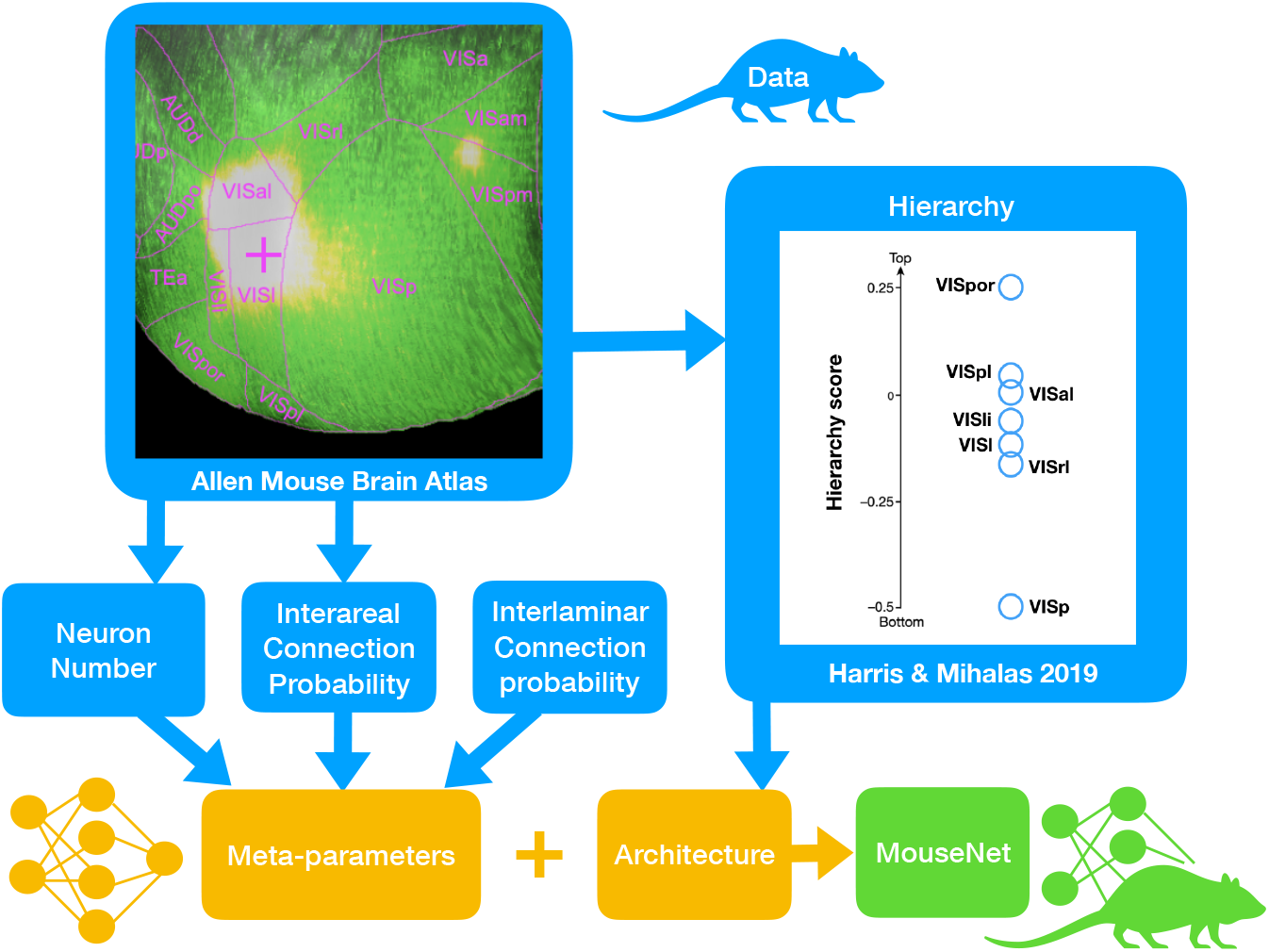
Modeling framework. Framework for constructing MouseNet from biological constraints on anatomy, via publicly available data from large-scale experiments. The CNN architecture is set by the analysis of hierarchy [23] on the Allen Mouse Brain Connectivity Atlas [20] (Image credit: Allen Institute); and the meta-parameters are mostly fixed by the combination of the 100-micrometer resolution interareal connectome [21] with detailed estimates of neuron density [35], and the statistics of connections between cortical layers from the literature [36–38].

### Network architecture

MouseNet spans the dorsal lateral geniculate nucleus (dLGN) and six visual areas (Fig 2A). Input to the network passes first through dLGN, and then to the primary visual area VISp. After VISp, the architecture branches into five parallel pathways, representing five secondary lateral visual areas: VISl (lateral visual area), VISal (anterolateral), VISpl (posterolateral), VISli (laterointermediate), and VISrl (rostrolateral). Finally, the output of VISp together with all five lateral visual areas provide input to VISpor (postrhinal). We include only the lateral areas because they are more associated to object recognition while the medial areas are more involved in multimodal integration [39]. The three-level architecture among the VIS areas was derived from an analysis of the hierarchy of mouse cortical and thalamic areas (Fig. 6e in [23]), which considered feedforward and feedback connection structures in each area. In this analysis, VISp was clearly low in the hierarchy, and VISpor was clearly high, but the other lateral visual areas had similar intermediate positions.

**Fig 2.**
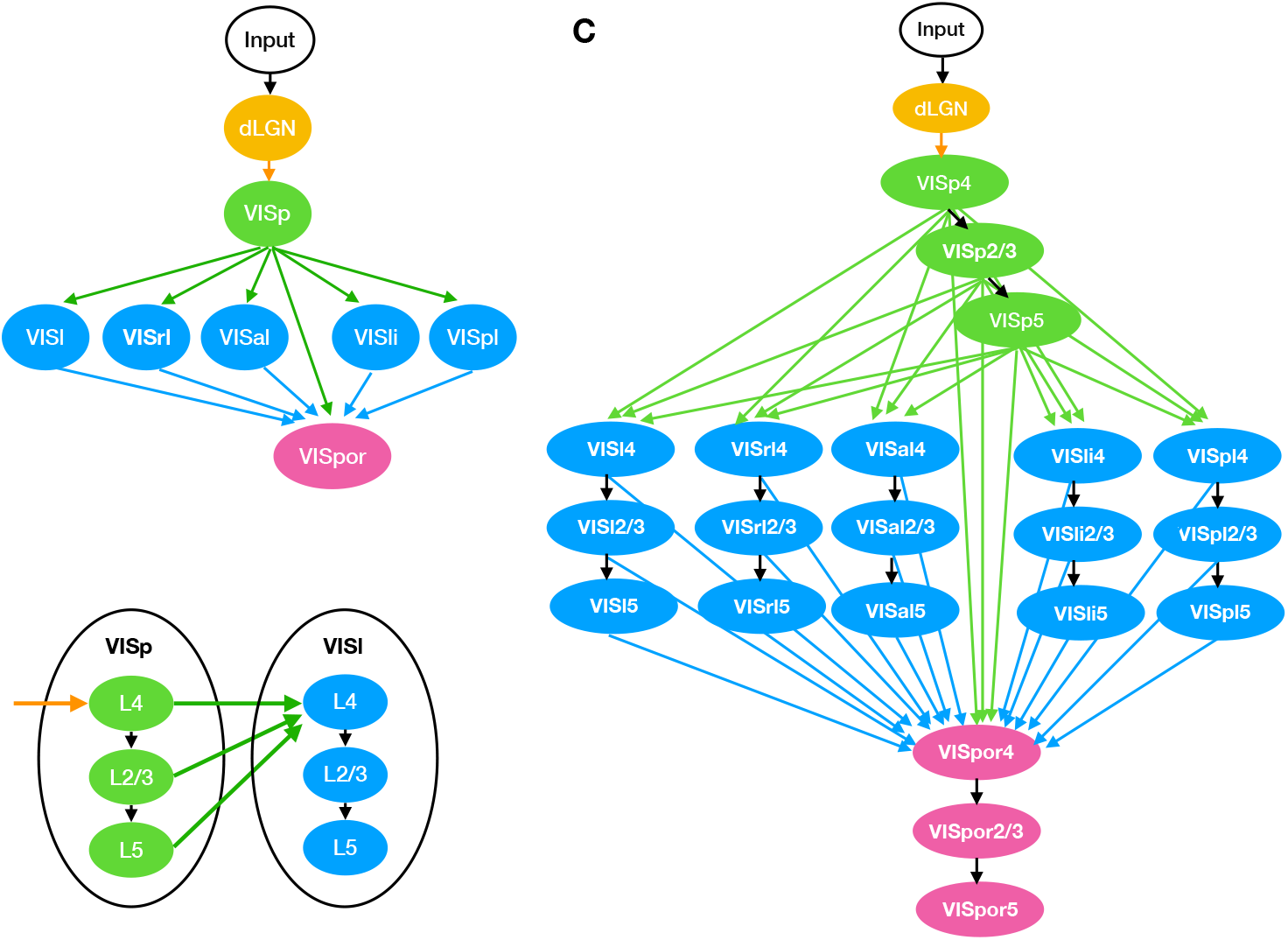
Illustration of MouseNet architecture. Only feedforward connections are included. (A) High-level organization of MouseNet, based on analysis of the hierarchy of lateral visual areas ([23]). (B) Connection patterns at the level of cortical layers. (C) Full MouseNet architecture.

In the MouseNet model, each VIS area is represented by three separate cortical layers: layer 4 (L4), layer 2/3 (L2/3) and layer 5 (L5). We call a specific cortical layer within a specific area a “region”. Here we only consider the feedforward pathway, thought in primate to drive responses within ≈ 100ms of stimulus presentation [2,8]. Following depictions of the canonical microcircuit (e.g. as summarized in Fig 5 in [40]), we consider the feedforward pathway to consist of laminar connections from L4 to L2/3, and from L2/3 to L5. Input from other areas feed into L4 and all of L4, L2/3 and L5 output to downstream areas, as shown in Fig 2B. This is consistent with broad connectivity among visual areas from each of these layers (Fig. 2f of [23]). Fig 2C shows the MouseNet architecture in full detail, including all 22 regions and associated connections.

### From architecture to convolutional neural net

Similar to the CALC model for the primate visual cortex by one of the authors [41], the general idea is to use convolution (Conv) operations to model the projections between different regions in the mouse visual cortex. Conv operations are linear combinations of many inputs, so they impose the assumption of linear synaptic integration. They are widely used in machine learning, because they run efficiently on graphical processing units, and they share parameters across the visual field, leading to reduced memory requirements and faster learning, relative to general linear maps.

Each connection from source brain region *i* to target brain region *j* is modelled with a Conv operation, Conv^*ij*^. The input to Conv^*ij*^ corresponds to the neural activities in source region *i*, and the output of Conv^*ij*^ drives neural activities in the target region *j*. For example, as shown in Fig 3A, the projection from Region 1 to Region 2 (Proj 1→2) is modeled by Conv^12^. The neural activities in Region 1 correspond to the input to Conv^12^, while the neural activities in Region 2 are a nonlinear function (ReLU, as shown in Fig 3C) of the output of Conv^12^. In MouseNet, L4 of all areas except VISp receive multiple converging inputs, similar to Region 4 in Fig 3A. In this case, each projection (Proj 2→4 and Proj 3→4) is modeled by a separate Conv layer (Conv^24^ and Conv^34^), and a nonlinear function (ReLU) is applied to the sum of the output from both of the Conv layers, to produce the neural activities in Region 4.

**Fig 3.**
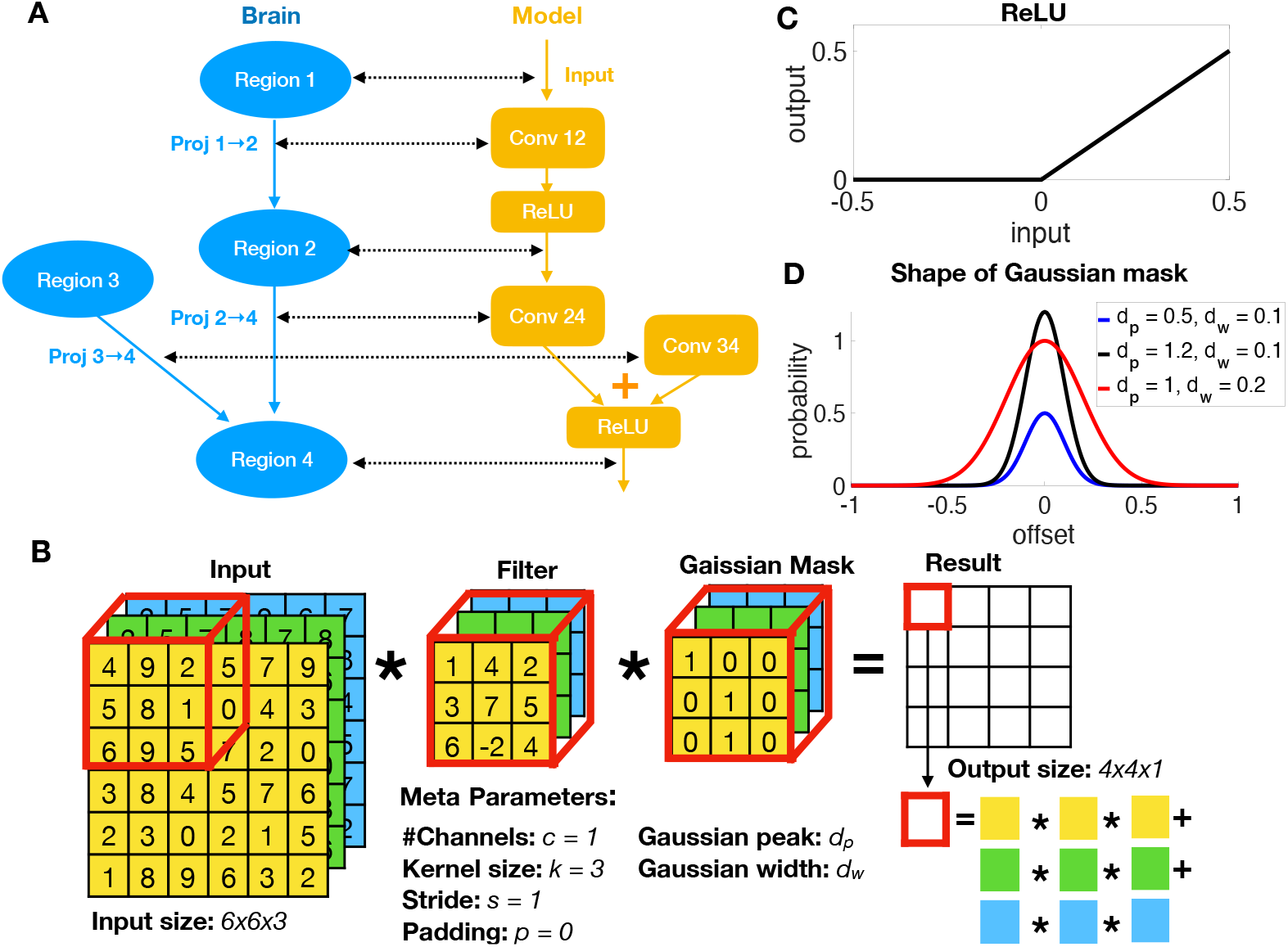
From mouse brain to CNN model. (A) From mouse brain hierarchy to CNN architecture. (B) An example of Conv operation with Gaussian mask. (C) ReLU operation in the CNN architecture. (D) The binary Gaussian mask is generated by a Gaussian shaped probability whose peak and width are meta-parameters.

### Finding meta-parameters consistent with mouse data

After fixing the architecture, we need to determine the meta-parameters for constructing the kernels for each Conv operation (Fig 3). The standard Conv operation is defined in terms of a four-dimensional kernel. The output of the kernel is a three-dimensional tensor of activations for region *j*, *A^j^*, which pass through element-wise ReLU nonlinearities to produce non-negative rates. Element 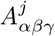 is the activation of the neuron at the *α^th^* vertical and *β^th^* horizontal position in the visual field, in the *γ^th^* channel (or feature map). The *γ^th^* channel of the activation tensor for region *j*, 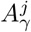, depends on inbound connections as,

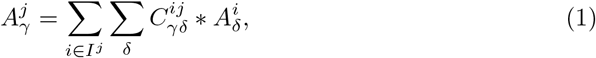

where *I^j^* is the set of regions that provide input to region j. Note that both 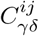 and 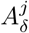 are two-dimensional, and they undergo standard two-dimensional convolution. The meta-parameters of kernel *C^ij^* are: number of input channels 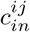, number of output channels 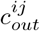, stride *s^ij^*, padding *p^ij^*, and finally kernel size *k^ij^*, i.e. the height and width (which are set equal) of 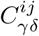. To make the connections realistically sparse, we add a binary Gaussian mask on the Conv operations, whose parameters are also estimated from data. See Fig 3B for an illustration of Conv operation with Gaussian mask. We constrain these meta-parameters with quantitative data whenever possible, and reasonable assumptions indicated by experimental observations otherwise, as indicated below.

#### Cortical population constraints

##### Assumptions about area output size

We set the horizontal and vertical resolution of the input (in pixels) based on mouse visual acuity. According to [42], the upper bound for visual acuity in mice is 0.5 cycles/degree, corresponding to a Nyquist sampling rate of 2 pixels/cycle x 0.5 cycles/degree = 1.0 pixel/degree. According to retinotopic map studies [43], V1 included a visual coverage range of ~ 60° in altitude and ~ 90° in azimuth, we further simplified this to square input size of 64 by 64 pixels.

The resolution of the other regions depends on both the resolution of the input, and strides of the connections. The stride of a connection is the sampling period with respect to its input. For example, a Conv with a stride of one samples every element of its input, whereas a Conv with a stride of two samples every other element (both horizontally and vertically), leading to output of half the size in each dimension. Because cortical neurons are not organized into discrete channels in the same way as convolutional network layers, there is no strong anatomical constraint on the stride. However, the mean stride has to be somewhat less than two; there are ten steps in the longest path through MouseNet, but if only six of them had a stride of two, the 64×64 input would be reduced to 1×1 in VISpor, with no remaining topography. Lacking strong constraints, for simplicity, we first attempted to set all the strides to one, but this left very few channels in some of the smaller regions (due to an interaction between channels and strides that we describe below). We therefore set the strides of the connections outbound from VISp to two, and others to one. The feature maps of dLGN and VISp were therefore 64×64 (the same as the input), and all others were 32×32.

Given the resolutions of the channels in each region, the numbers of channels are constrained by the number of neurons. Specifically, Let *n^i^* be the number of neurons in region *i* and 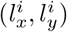 be the size of the output in area *i*, then the number of channels in area *i* is determined by 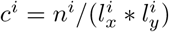.

##### Estimating number of neurons in each area from data

We only model the excitatory neural population in our model, consistent with the fact that all neurons in the model project to other visual areas. In fact, neurons in convolutional networks are neither excitatory or inhibitory, but have both positive and negative output weights. However, past modelling work [44, 45] has shown that such mixed-weight projections can be transformed so that the original neurons are all excitatory, and an additional population of inhibitory neurons recovers the functional effects of the original weights.

According to [46], the estimated number of excitatory neurons in dLGN is 21200. For VISp, VISal, VISl, VISpl, we use estimated density for excitatory neurons given by [35]^1^, which is summarized in Tabel 1. Note that we use neuron density instead of counts to get a more stable estimation of number of neurons out of different versions of brain parcellations. For the remaining areas VISrl, VISli and VISpor, we approximate their density by taking the average across the above four areas with separated cortical layers.

**Table 1.** Exitatory population density [mm^-3^] [35].

|  | L4 | L2/3 | L5 |
| --- | --- | --- | --- |
| VISp | 106114.7 | 86668.2 | 86643.4 |
| VISal | 93176.9 | 79070.6 | 78540.9 |
| VISl | 86559.9 | 73937.9 | 66215.6 |
| VISpl | 106783.0 | 87368.3 | 82538.1 |
| Average | 98158.6 | 81761.1 | 78484.5 |

Combined with the number of 10μm voxels counted in the Allen Mouse Brain Common Coordinate Framework (CCFv3) [47] (Table 2), we summarize the estimated number for all the regions in our model in Table 3.

**Table 2.** Number of 10μm voxels in each region.

|  | L4 | L2/3 | L5 |
| --- | --- | --- | --- |
| VISp | 1023640 | 1999040 | 1552688 |
| VISal | 104152 | 199314 | 202942 |
| VISl | 179084 | 301588 | 314522 |
| VISpl | 36638 | 205150 | 242812 |
| VISrl | 146294 | 276390 | 244294 |
| VISli | 57256 | 117252 | 147946 |
| VISpor | 60632 | 373972 | 385168 |

**Table 3.**
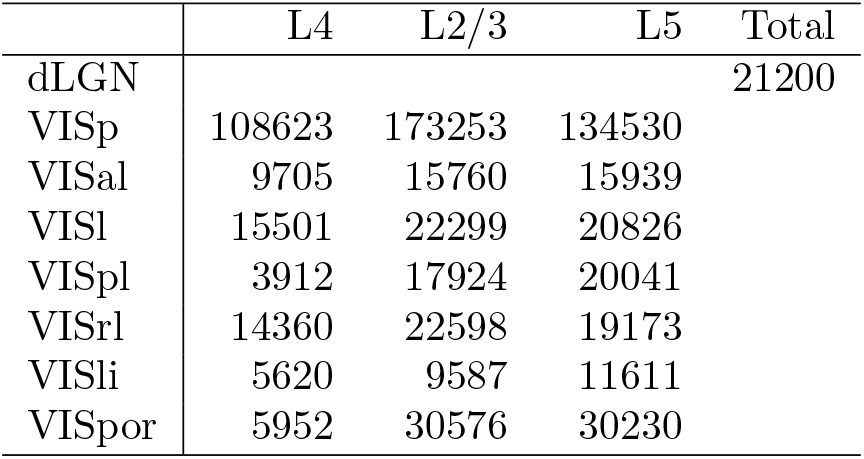
Estimated number of exitatory neurons in each region.

|  | L4 | L2/3 | L5 | Total |
| --- | --- | --- | --- | --- |
| dLGN |  |  |  | 21200 |
| VISp | 108623 | 173253 | 134530 |  |
| VISal | 9705 | 15760 | 15939 |  |
| VISl | 15501 | 22299 | 20826 |  |
| VISpl | 3912 | 17924 | 20041 |  |
| VISrl | 14360 | 22598 | 19173 |  |
| VISli | 5620 | 9587 | 11611 |  |
| VISpor | 5952 | 30576 | 30230 |  |

#### Cortical connection constraints

Neurons tend to receive relatively dense inputs from other neurons that are above or below them, in other cortical layers, and the connection density decreases with increasing horizontal distance. Similarly, inputs from other cortical areas tend to have a point of highest density, with smoothly decreasing density around that point. We approximate such connection-density profiles with two-dimensional Gaussian functions. Specifically, the fan-in connection probability from source region *i* to target region *j* at position (*x, y*) (position offset from center in μm) is modeled as,

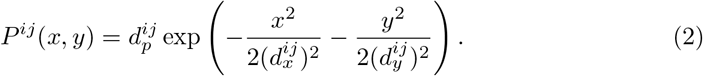

where 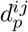 is the peak probability at the center and 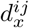 and 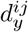 are the widths in the *x* and *y* directions. For simplicity, we assume 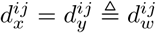 and let 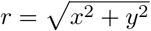 denote the offset from the center of the source layer, the above equation then simplifies to,

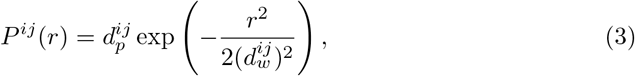

where 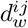 (μm) is the Gaussian width.

Both 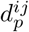 and 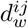 are estimated from mouse data. The parameters for interlaminar connections are estimated from statistics of connections between cortical layers in paired electrode studies (Section Estimating 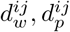 for interlaminar connections), and the parameters for interareal connections are estimated from the mesoscale mouse connectome (Section Estimating 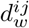 and 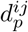 for interareal connections).

#### Conv layer with Gaussian mask

To translate our Gaussian models of connection density into network meta-parameters, we apply a binary mask to the weights of the Conv layers (Fig 3B). To do that, we first change the unit of 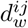 in Eq. 3 from micrometers to source area-dependent “pixels” (unit of output size of source area *i*) by multiplying it with 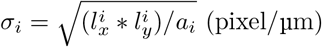, where *a_i_* denotes the surface size of area *i*, estimated from the voxel model (See Estimating 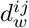 and 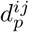 for interareal connections). We then have,

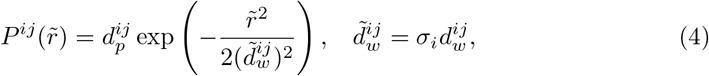

where both 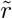 and 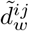 are in the “pixel” unit. The kernel size of the Conv layer is set to be 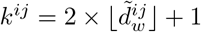, with padding calculated by 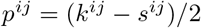, where *s^ij^* is the stride of the Conv layer.During initialization, a mask containing zeros and ones is generated for each Conv layer, with size 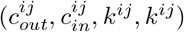. The probability of each element being one is 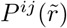, where 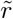 (pixel) is the offset from the center of mask. The weights of the Conv layer are then multiplied by the mask. This gives the connections realistic densities (or sparsities), with realistic spatial profiles.

#### Estimating 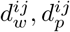 for interlaminar connections

For the interlaminar connections, we estimate the Gaussian width 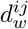 from multiple experimental resources. Firstly, from Table 3 in [37], we get the estimation of 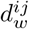 to be 114 micrometers for functional connections between pairs of L4 pyramidal cells in mouse auditory cortex. Secondly, manually extracted from [38] Fig 8B, we obtain the variation of the Gaussian width depending on source and target layer from cat V1. Finally, we use this variation to scale the L4 to L4 width of 114 μm to other layers in the mouse cortex. We summarize the Gaussian widths from cat cortex, along with corresponding scaled estimates for mouse cortex, in Table 4. Note that in the current model, we only use the values for connections from L4 to L2/3 and from L2/3 to L5 (Fig 2B).

**Table 4.** Estimated Gaussian width 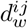 for interlaminar excitatory connections. The values outside of the parenthesis are extracted from [38]; the values inside the parenthesis are scaled to mouse cortex, using the width 114 μm for L4-to-L4 connections in mouse auditory cortex [37]. Units are micrometers (μm).

|  |  | Target |  |  |
| --- | --- | --- | --- | --- |
|  |  | L2/3 (scaled) | L4 (scaled) | L5 (scaled) |
| Source | L2/3 | 225 (142.5) | 50 (31.67) | 100 (63.33) |
|  | L4 | 220 (139.33) | 180 (114) | 140 (88.67) |
|  | L5 | 150 (95) | 100 (63.33) | 210 (133) |

To estimate the Gaussian peak probability 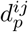, we first collect the connection probability between excitatory populations offset at 75 micrometer 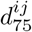 (Fig. 4A in [36]). We then get the peak probability 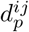 by the relation

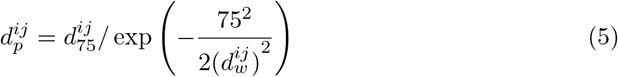

**Fig 4.**
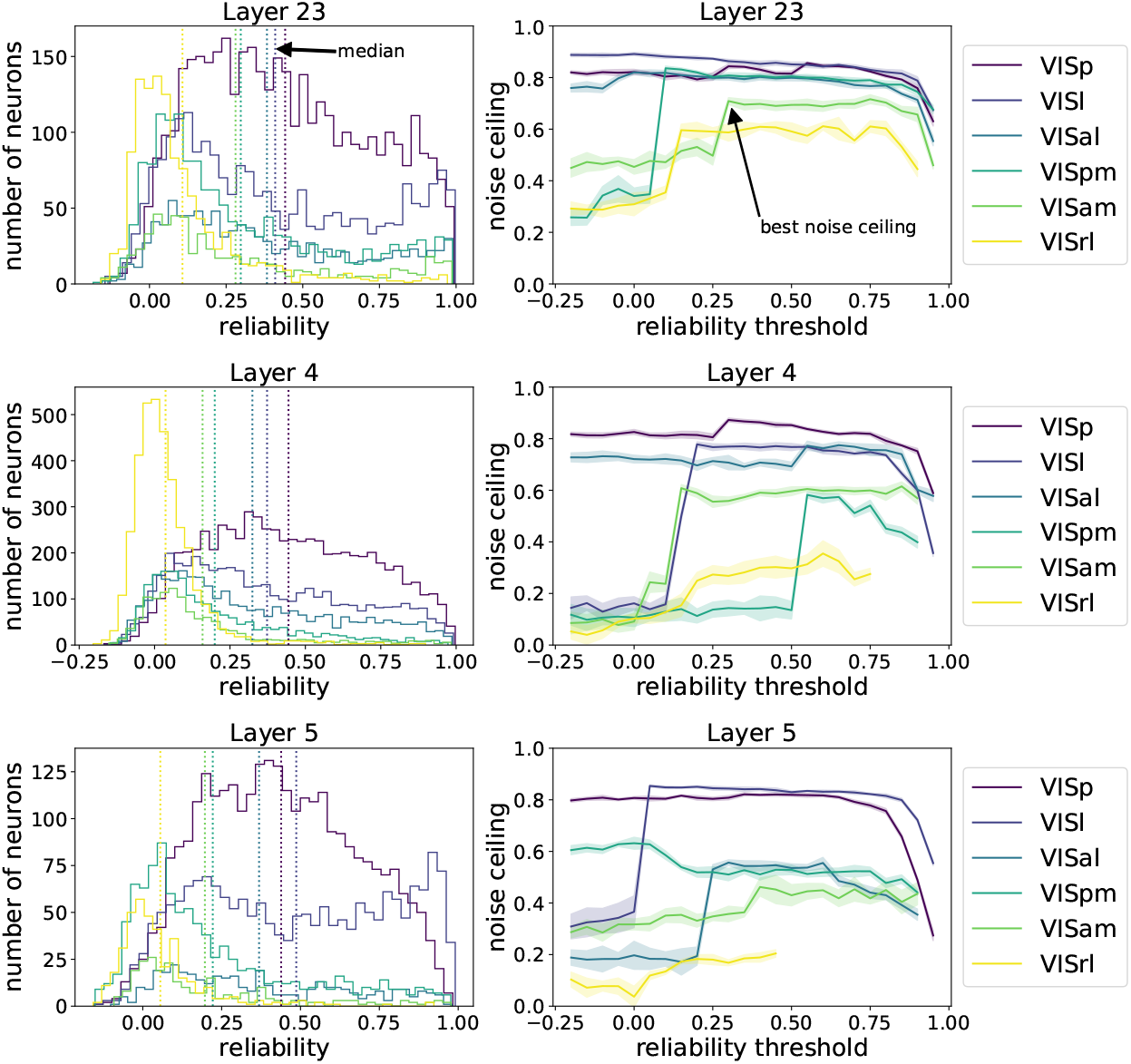
Selecting reliable neurons improves noise ceilings. (Left) Reliability distribution of neural populations. Each row shows all the brain areas at a specific cortical layer. The dotted lines indicate the median reliability of each neural population. (Right) The noise ceilings change with variation of the threshold for selecting reliable neurons. The higher the threshold, the fewer neurons are selected. For some populations, selecting a certain portion of reliable neurons gives best noise ceiling. Error bars are from different draws of non-overlapping trials.

We summarize the probability at 75 micrometers 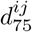 along with the peak probability 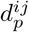 in Table 5.

**Table 5.** The connection probability between excitatory populations offset at 75 micrometer. 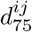 The numbers are from Fig 4A in [36]). The calculated Gaussian peak probability 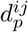 are given in parenthesis.

|  |  | Target |  |  |
| --- | --- | --- | --- | --- |
|  |  | L2/3 (peak) | L4 (peak) | L5 (peak) |
| Source | L2/3 | 0.160 (0.184) | 0.016 (0.264) | 0.083 (0.167) |
|  | L4 | 0.140 (0.162) | 0.243 (0.302) | 0.104 (0.149) |
|  | L5 | 0.021 (0.029) | 0.007 (0.014) | 0.116 (0.136) |

#### Estimating 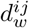 and 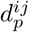 for interareal connections

To estimate interareal connection strengths and spatial profiles, we use the mesoscale model of the mouse connectome [21, 22]. This model estimates connection strengths between 100 micrometer resolution voxels, based on 428 individual anterograde tracing experiments mapping fluorescent labeled neuronal projections in wild type C57BL/6J mice.

##### Flat map

The voxel model is in 3 dimensional space. For the purpose of our analysis, we need to map the 3 dimensional structure into 2 dimensions. First, we fit the visual area positions by a sphere with center 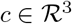 and radius *r*. Each position 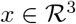 is then mapped to 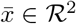 with relation

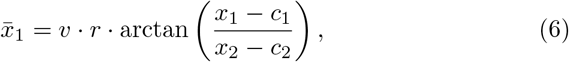

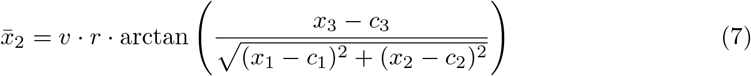

where *v* = 100μm is the size of the voxel.

##### Area size

Approximations of the surface area for each brain region are needed to convert the widths of connection profiles (see Conv layer with Gaussian mask) from voxels in the mesoscale model to convolutional-layer pixels in MouseNet. For this purpose, each region’s surface area size is approximated by the area of a convex hull of its mapped two-dimensional positions. These estimates are summarized in Table 6.

**Table 6.** Area size (mm^2^) estimated from the voxel model.

|  | VISp | VISal | VISl | VISli | VISpl | VISrl | VISpor |
| --- | --- | --- | --- | --- | --- | --- | --- |
| L4 | 4.3271 | 0.4909 | 0.8793 | 0.3355 | 0.2865 | 0.6182 | 0.5264 |
| L2/3 | 4.7406 | 0.5477 | 0.9279 | 0.4356 | 0.6659 | 0.6980 | 1.3937 |
| L5 | 4.2511 | 0.4972 | 0.8651 | 0.4039 | 0.6785 | 0.6748 | 1.2445 |

##### Estimating 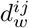

For each connection from source region *i* to target region *j*, we estimate 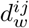 from the mesoscale model. The first step is to estimate the widths of connections to individual voxels in *j*. The incoming width 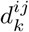 for target voxel *k* in *j* is estimated by the standard deviation of the connection strength about its center of mass. Specifically, 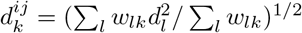, where *l* indexes the voxels in source region *j*, *w_lk_* is the connection weight between source and target voxels *l* and *k* in the mesoscale model, and *d_l_* is the distance from voxel *l* to the center of mass of these connection weights. We then estimate 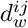 as the average of these widths over the voxels in *j*. We omit from this average any target voxels that have multi-modal input profiles. This procedure provides an upper bound for 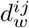, because a target voxel may include multiple neurons with partially overlapping input areas.

##### Estimating 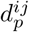

The mesoscale model provides estimates of relative densities of connections between pairs of voxels. But an additional factor is needed to convert these relative densities into neuron-to-neuron connection probabilities. For this purpose, we assumed that each neuron received inputs from 1000 neurons in other areas (we call this number the extrinsic in-degree, *e*). This is on the order of the estimate from Fig S9 M in [48]. Given this assumption, we calculated 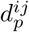 by the relation,

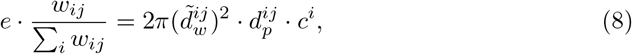

where *w_ij_* is the connection strength from source area *i* to target area *j*, estimated from integrating the connection weights of the corresponding areas in the mesoscale model. The estimated values for 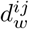 and 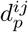 are given in Table 12 in S1 Table.

#### Conv kernel size for dLGN

The above methods allowed us to set kernel sizes for intracortical connections, but not subcortical ones. We set the kernel sizes for inputs to dLGN and VISp L4 according to receptive field sizes in these regions. Receptive fields are about 9 degrees in dLGN and 11 degrees in VISp [49]. As mentioned in Section Cortical population constraints, mouse visual acuity is approximately 1 pixel/degree, therefore we set kernel size of the connection from input to dLGN to 9×9. We then set the kernel size of the connection from dLGN to VISp to 3×3, such that the receptive field size for VISp is 11×11 pixels.

#### Summary tables

In Table 7, we summarize the calculated number of channels in each area (in parenthesis) and the kernel size for each Conv layer.

**Table 7.**
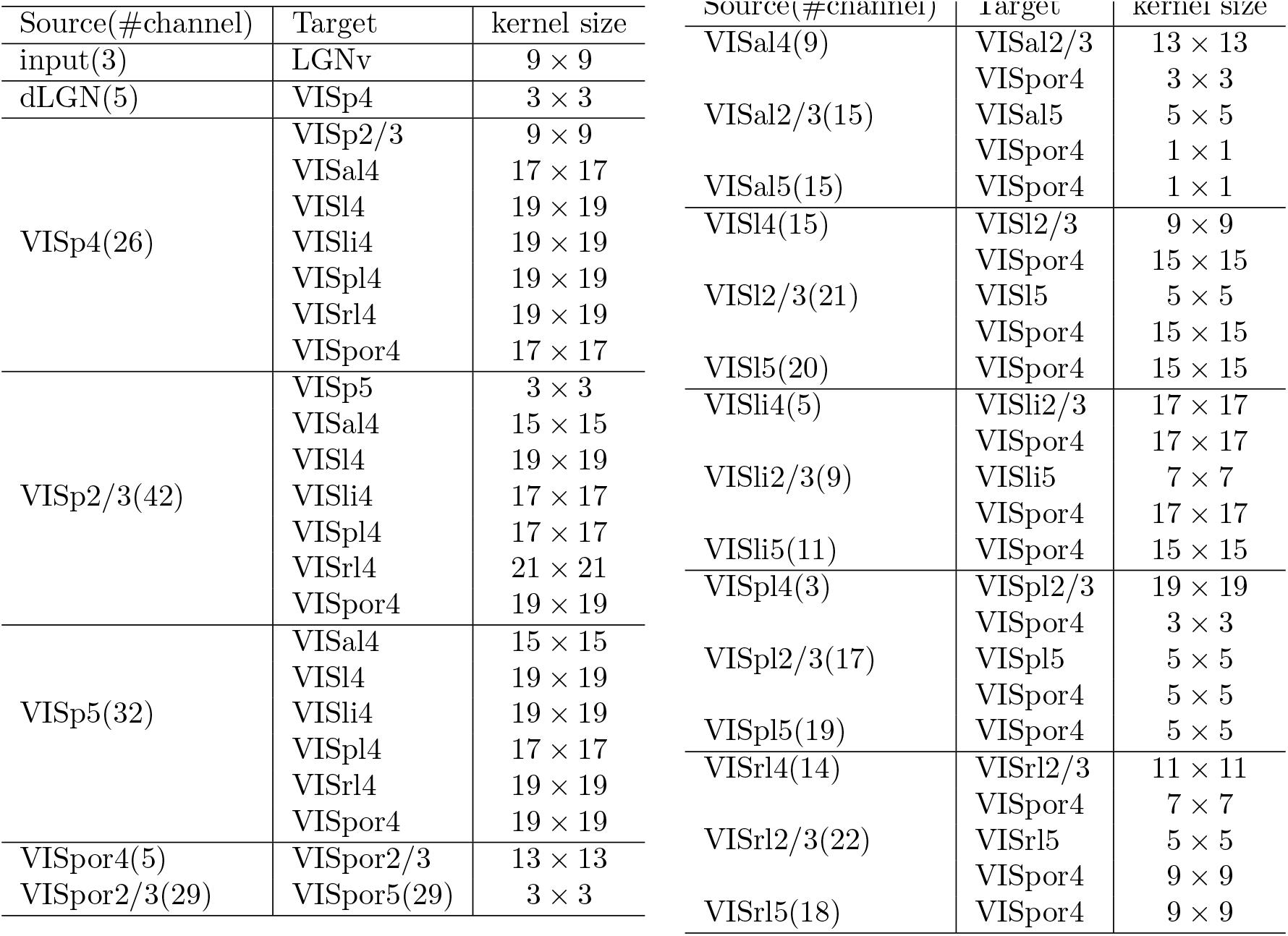
The calculated meta-parameters for the Conv layers.

The parameters used in the model based on biological sources and assumptions are summarized in Table 8 and the formulae for calculating the Conv layer meta-parameters are sumarized in Table 9.

**Table 8.** Parameters from data or assumptions.

| Notation | CNN parameter | Biological source or assumptions |
| --- | --- | --- |
| $n^i$ | Number of neurons in area $i$ | Based on [35] combined with the voxel model [21] |
| $a_i$ | Two dimensional area size for area $i$ | Estimated from voxel model data [21] |
| $e$ | Total fan-in connections for all areas | Set to be 1000 based on [48] |
| $(l_x^0, l_y^0)$ | Input size to the model | Set to be 64x64 based on mouse visual acuity [42] |
| $(l_x^i, l_y^i)$ | Output size of area $i$ | Set to be 64x64 up to VISp, 32x32 after VISp (Assumption) |
| $d_w^{ij}$ | Gaussian width (interlaminar) | Estimated from mouse [37] and cat [38] cortical properties |
|  | Gaussian width (interareal) | Estimated from voxel model [21] |
| $d_p^{ij}$ | Gaussian peak (interlaminar) | Based on statistics of connections in paired electrode studies [36] |
|  | Gaussian peak (interareal) | Estimated from voxel model [21] |

**Table 9.**
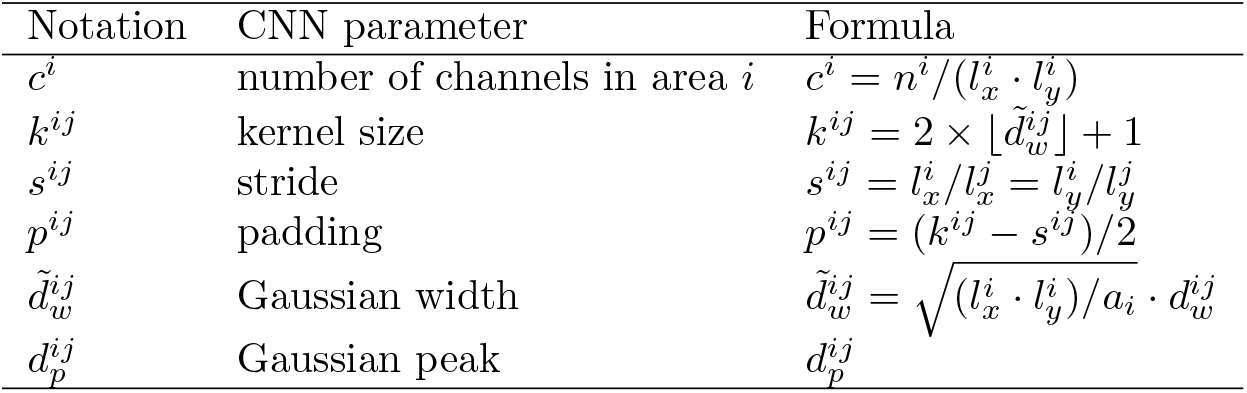
Meta-parameters for Conv layer connecting source area *i* to target area *j*.

## Results

In this section, we use a well established image classification task as a working example to demonstrate the usage of the CNN MouseNet and to derive novel findings relating architecturally constrained CNNs and the mouse brain. We first assess the computational performance of this mouse-architecture network on an image classification task. Then, through systematic comparisons with the large scale Allen Brain Observatory dataset, we show how MouseNet can be used to probe the effect of a CNN’s specific task training and architecture on its similarities and differences with responses in the biological brain.

### Implementation of MouseNet

To enable training of MouseNet on a standard image classification task, we implemented the MouseNet structure in PyTorch [50]. Each Conv layer was followed by a batch normalization layer and a ReLU non-linearity. For regions such as VISpor L4 that receive input from multiple Conv layers, the outputs of the Conv layers are summed before being fed into the batch normalization layer and ReLU non-linearity.

To train the MouseNet model on an image classification task, we added a simple classifier. Specifically, in order to include the final processing output from each individual area such that the information is not bottlenecked by the relatively small VISpor area, we took the L5 output from all seven areas and reduce them to 4×4 by an average pooling layer. We then flattened, concatenated, and fed this to a linear fully-connected layer, which reduced the dimension to the number of classes of the task. The outputs were then transformed to probabilities by the softmax function, and the cross-entropy loss of the predicted probabilities (determined from the categorical distribution where individual class probabilities are from the output of the softmax) relative to the ground truth labels was used to train on the image classification task.

### Computational Performance of MouseNet on image classification

We trained MouseNet end-to-end using stochastic gradient decent with momentum, adapting the training script from the imagenet example script from the PyTorch examples github repository^2^. Full training details and scripts are available on the MouseNet github repo: https://github.com/mabuice/Mouse_CNN.

We first found that MouseNet achieved above 90% validation accuracy on CIFAR10 [31], a simple classification of 32×32 images into 10 categories. Interestingly, this is close to state of the art performance of modern networks, suggesting that mouse sized networks are fully capable of performing this simple task.

We then moved to the more challenging image classification benchmark of ImageNet [51], which requires classification of higher resolution images into 1000 categories. We found that, even for input images downsampled to a resolution of (64×64), MouseNet can still be trained to perform above 37% top-1 validation accuracy on ImageNet [51]. Below, we contrast representations in MouseNet to those in VGG16 trained with the same downsampled input size (64×64), which achieved above 60% top-1 validation accuracy on ImageNet. We contrast the number of parameters in MouseNet and VGG16 in Table 10. Note that the number of parameters of MouseNet Conv layers without the Gaussian masks is about 14% of that for VGG16, while the number parameters of MouseNet Conv layers with Gaussian masks is less than 1% of that for VGG16. Our simulation results are all based on MouseNet models with Gaussian masks.

**Table 10.** Number of parameters for MouseNet and VGG16 for 1000-class ImageNet classification task.

|  | Conv layers | Conv with mask | Classifier |
| --- | --- | --- | --- |
| VGG16 | 14.7M | 14.7M | 123M |
| MouseNet | 2.1M | 87K | 2.3M |

### The Effects of Task Training on Functional Properties

To examine the effect of the image classification task training on the functional similarity of the MouseNet and the biological mouse brain, we make use of the large-scale, publicly available Allen Brain Observatory dataset [26]. We study representational similarity of MouseNet and the biological mouse brain across a set of natural images, along with the basic functional properties of sparsity and orientation selectivity.

#### The Allen Brain Observatory data set

The Allen Brain Observatory data set is a large-scale standardized *in vivo* survey of physiological activity in the mouse visual cortex, featuring representations of visually evoked calcium responses from GCaMP6f-expressing neurons. In this work, we use the population neural responses to a set of 118 natural image stimuli, each presented 50 times. The images were presented for 250ms each, with no inter-image delay or intervening “gray” image. The neural responses we use are events detected from fluorescence traces using an L0 regularized deconvolution algorithm, which deconvolves pointwise events assuming a linear calcium response for each event and penalizes the total number of events included in the trace. Full information about the experiment is and database given in [26].

The Allen Brain Observatory includes data from six different brain areas, namely VISp, VISal, VISl, VISpm, VISam and VISrl. The number of neurons in the dataset, for each of the regions we use, is summarized in Table 11.

**Table 11.** Number of neurons recorded from each mouse brain region.

|  | VISp | VISal | VISl | VISpm | VISam | VISrl |
| --- | --- | --- | --- | --- | --- | --- |
| Total | 14173 | 4396 | 8748 | 4771 | 2040 | 5189 |
| L2/3 | 4079 | 1042 | 2259 | 1544 | 610 | 1168 |
| L4 | 6735 | 2967 | 4163 | 1905 | 1179 | 3626 |
| L5 | 3003 | 387 | 1874 | 973 | 251 | 395 |

#### The Similarity of Similairy Matrices metric (SSM)

To compare functional similarity betweeen two representations – in MouseNet, and in the biological mouse brain – of a set of images, we use the Similarity of Similarity matrices (SSM) [27, 32] metric. We begin with a matrix of neural activities, in which each row contains the population activities for a certain image. We calculate the Pearson correlation coefficient between every pair of rows within one representation matrix, to form an *n* by *n* “similarity matrix” for this representation, where each entry describes the similarity of the population response to a pair of images. Next, to compare two similarity matrices, we flatten the matrices to vectors and compute the Spearman rank correlation between these vectors. Like the Pearson correlation coefficient, the rank correlation lies in the range [-1,1] indicating how similar (close to 1) or dissimilar (close to −1) the two representations are. Rather than examining one neuron at a time [52, 53], this metric compares representations based on activities of the whole populations of artificial and biological neurons, revealing functional similarity at the population level. Another choice of such population similarity metrics is Singular Vector Canonical Correlation Analysis (SVCCA) [27,54]. An excellent review of such similarity metrics and their properties can be found in [55].

Following the procedures in [27], we construct the representation matrix for a certain mouse visual cortex region by taking the trial-averaged mean responses of the neurons in the 250ms during the image presentation. Activities of neurons in different experiments for the same brain area are grouped together to construct the representation matrix, whose dimension is number of images by number of neurons. The representation matrices for MouseNet layers are obtained from feeding the same set of 118 images (resized to 64×64) to MouseNet and collecting all the activations from a certain layer of the model.

#### Neural reliability and SSM noise ceiling

We next compute the SSM noise ceiling from the Allen Brain Observatory data. We use split half reliability to quantify the reliability of a single neuron from the Allen Brain Observatory. This is done by separating the 50 trials into two non-overlapping 25 trial sets, and taking the correlation of trial-averaged responses between the two. We make ten random splits, and take the mean of the ten correlations to represent the reliability of each neuron. The reliability distributions of the neural populations are shown in Fig 4 (left). VISp, VISl and VISal are most reliable areas and VISpm, VISam and VISrl are less reliable areas.

To estimate the noise ceiling of the SSM metric, we compare the mouse data representation matrices with themselves. Specifically, we split the 50 trials in the dataset into two non-overlapping sets and calculate the trial averaged representation matrices for each set. The SSM between these two representation matrices are the noise ceiling of the SSM metric. Multiple splits of the dataset are computed for estimating the mean and standard deviation of the noise ceilings.

To examine how the noise ceiling changes with the reliability of the neural population, we calculate the noise ceilings by selecting neurons that surpass different levels of thresholds, as shown in Fig 4 (right). We see that for some regions, if we select a group of neurons using a certain reliability threshold, the noise ceiling becomes higher than without this selection. We summarize the reliability and best noise ceiling for each area in Fig 5. In this paper, we will concentrate our discussions on the most reliable areas, VISp, VISl and VISal, which are included in the MouseNet model. We will use the best noise ceiling to compare with the models.

**Fig 5.**
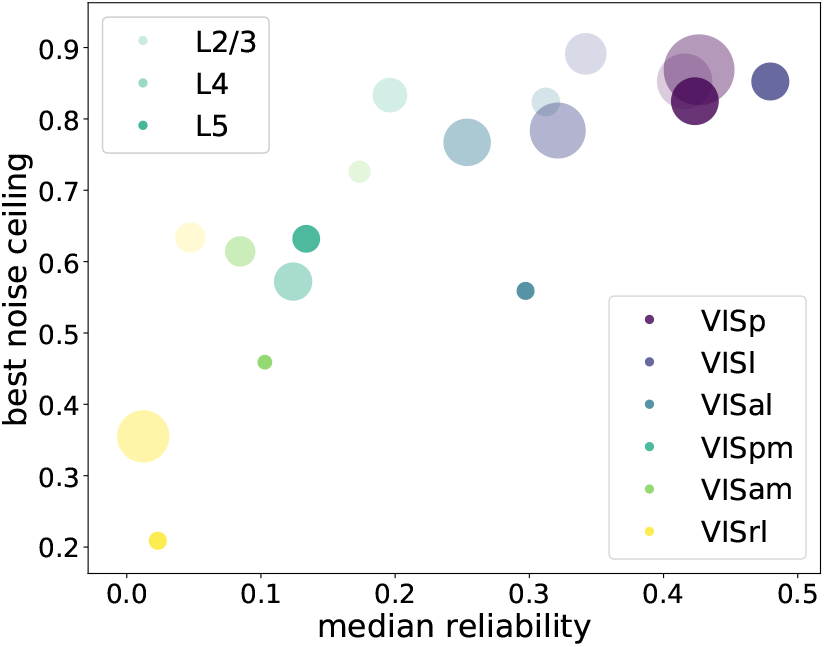
Summary plot of median reliability and best noise ceiling for each brain area. Each color represents a different brain area, and shades from light to dark indicate different cortical layers L2/3, L4 and L5. The circle size is proportional to the size of the population in the dataset.

#### Task training improves the similarity between MouseNet and the Allen Brain Observatory

To examine the effect of training to perform an image classification task on the functional similarity of MouseNet to the brain, we compute the SSM value between each layer of MouseNet with data from a brain region recorded in the Allen Brain Observatory. To account for the randomness due to initialization, we train four instances of MouseNet on ImageNet starting with different weights and look at their mean statistics. Fig. 6 shows the SSM values between each of the MouseNet layers with data from L2/3 of VISp, VISl and VISal. Layers 4 and 5 are shown in Fig. 11. The first important observation is that regions in the model do not necessarily best match to the corresponding functional area recorded in the Allen Brain Observatory. We see that for layer 2/3, area VISp in the Allen Brain Observatory, five different model areas show significant change in SSM value from the untrained model. In the following, we will add prefix “m” in front of the modeled areas from the MouseNet to contrast with the ones from the real brain. One of these is an early layer, mVISp5, while the others are in the parallel pathway portion of the architecture. Of the others, mVISl4 shows an increase in similarity with VISp_layer23, while three other model regions show a decrease in similarity. For the other two regions in Figure 6, mVISp5 shows a significant increase in similarity. For VISl_layer23, there are six other model regions that all show an increase in similarity. These statements hold specifically when comparing model regions to each other for the same area in the Brain Observatory. Comparing areas of the Brain Observatory to each other requires a different adjustment for the number of comparison (see black vs. red stars in Figure 6). These results are consistent with the idea from Shi, et al [27] that VISp is a lower order area than VISl and VISal (VISp maps to lower “pseudo-depth” in comparing to a CNN than both VISl and VISal). Layers 4 and 5 show results that are similar, but not identical to, layer 2/3. (Fig. 11). VISal in Layer 4 and VISl and VISal in Layer 5 show improved similarity after training for many of the mVISp model regions. Similarly, VISp in layer 4 and 5 shows decreased similarity after training in some of regions in the parallel portion of the architecture.

**Fig 6.**
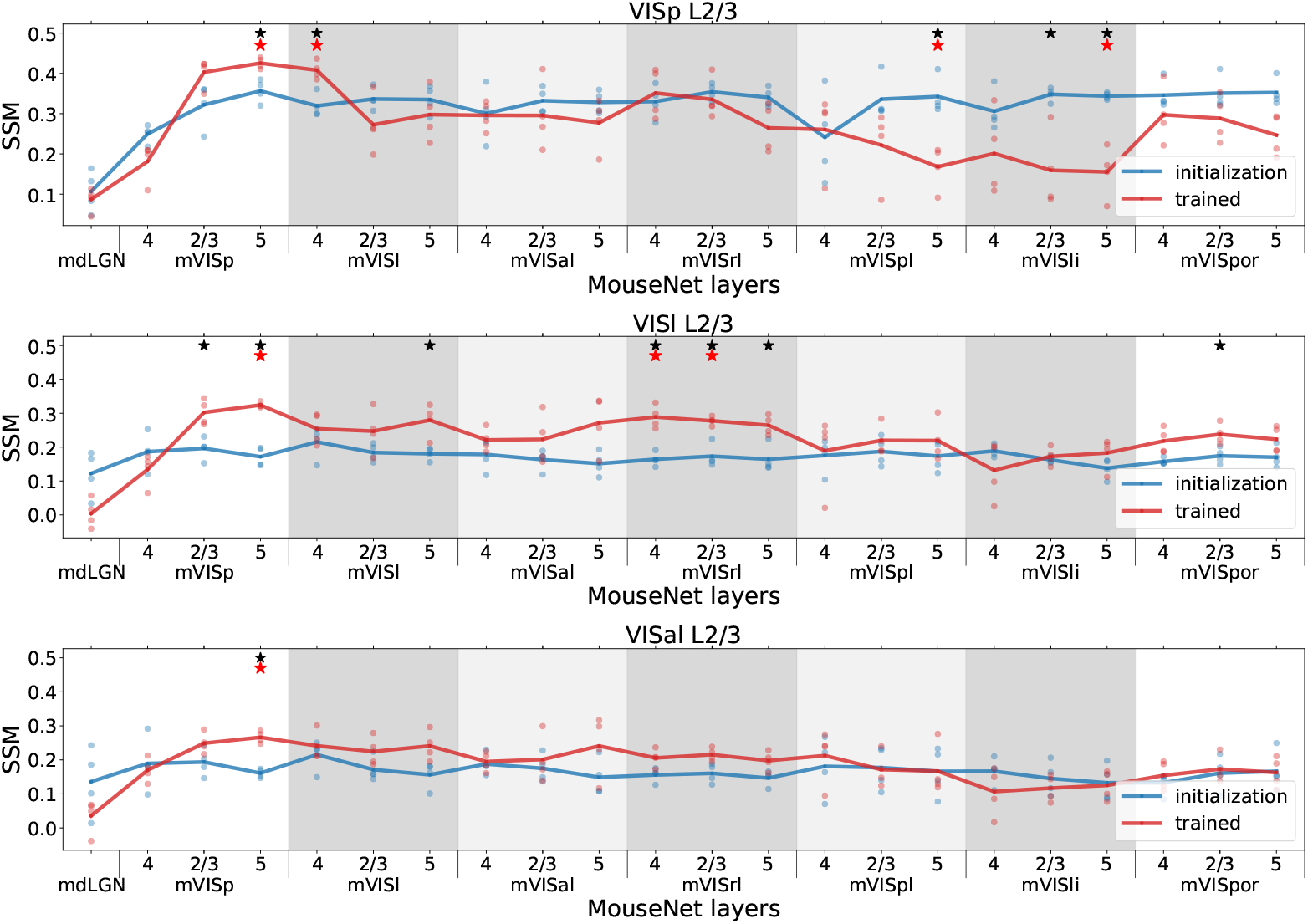
SSM between mouse data in VISp(top)/VISl(middle)/VISal(bottom) L2/3 and all layers in the MouseNet before (blue) and after training (red). Each line corresponds to the mean of four different MouseNet instances trained from different initialization weights (dots). The x axis includes all the layers in the model in a serial way. The five parallel secondary visual area pathways in the model are in shaded grey background. Black stars denote the the pvalues of two-sample t-test with Benjamini/Hochberg correction of 22 comparisons within one brain area is less than 0.05; Red stars denote the pvalues of two-sample t-test with Benjamini/Hochberg correction of all 9×22 comparisons across all 9 brain areas is less than 0.05.

Note that, although training on ImageNet improves the corresponding level of model regions’ similarity to the brain, the highest SSM value does not always occur in the model layer corresponding to the specific region considered in the Brain Observatory. For example, the SSM value for mVISp regions are higher than the mVISl regions when comparing to the brain area VISl L2/3. This is possibly because the visual areas are more similar to each other than they are to the MouseNet regions (see Table 13 in S1 Table for the SSM values between the brain areas themselves), such that improving the similarity to one brain region can possibly lead to improving the similarity to some other regions. Nevertheless, by looking at all the layers globally, we see that for the earliest visual area VISp, the ImageNet classification training promotes the SSM values of the mVISp layers in the MouseNet while suppress the values for the later layers; whereas for secondary visual areas VISl, the training promotes both earlier layers and later layers in the parallel pathways, suggesting a higher place in the functional hierarchy (cf. with the results of [27]).

#### Higher task performance on image classification does not guarantee higher similarity to the mouse brain

To examine how performance on the ImageNet classification task affects the functional similarity to the brain, we contrast the SSM values for MouseNet with another network that can perform this task, the VGG16 network discussed above. We use the same input resolution, on the same task (see Section Computational Performance of MouseNet on image classification). Similarly as for MouseNet, we calculate the SSM values between each layer in VGG16 and the regions in the mouse visual cortex. VGG16 does not have a “corresponding layer” for each region; we choose the VGG layer that has the highest SSM with each mouse brain region. For this comparison, we do the same for MouseNet, so that for each region, we compare this ‘best layer’ SSM value with the best layer SSM value for MouseNet.

The best layer’s SSM values for both VGG16 and MouseNet, for each mouse cortical layer in VISp, VISl, and VISal, are summarized in Fig 7. As we can see in the figure, although VGG16 has much higher performance on the ImageNet task (about 60% vs 40%), it does not have much higher SSM values to the brain for most regions. The saturation of functional similarity between the brain and models in terms of image classification performance is also observed in primates, albeit at a much higher performance level [56].

**Fig 7.**
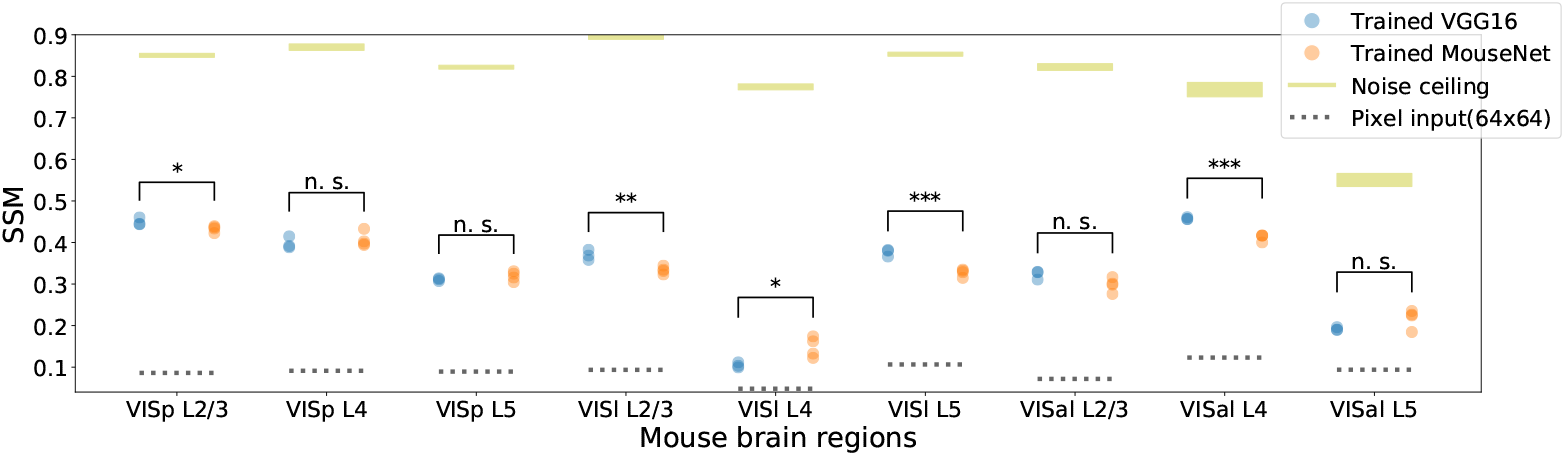
SSM between best layer in trained VGG16/MouseNet and mouse brain regions. The plot shows results of 3 instances of VGG16 (with validation accuracy 60.46, 60.72, 60.93) and 4 instances of MouseNet (with validation accuracy 37.46, 37.95, 37.52, 37.49) trained from different initialization weights. Yellow lines denote the best noise ceiling; their widths are standard deviations calculated from multiple draws of non-overlapping trials as in Fig.4. Dotted black lines are the SSM values between the 64×64 pixel input and the corresponding regions. Black stars denote the statistical significance of two-sample t-test between the mean of the trained VGG16 and the trained MouseNet instances (one star: p < 0.05, two stars: p < 0.01, three stars: p < 0.001).

To further grasp the limited relationship between a model’s task performance and its functional similarity to the mouse brain, we look at how the models’ functional similarity to brain data changes during training. As shown in Fig 8, the highest SSM values between a model neural network and the mouse brain areas are not necessarily achieved by the best performing models, rather at early or intermediate points during the training process. See Fig.12 in S1 Fig for more instances of MouseNet during training, also showing this effect. These results show that optimizing performance on this particular task, at least beyond an early or intermediate level of performance, does not necessarily improve the model’s similarity to the biological brain. If the approach of building models for neural responses via task training of artificial networks is broadly correct, then we take this as an indication that ImageNet is not the correct task to consider for the representations in the mouse brain.

**Fig 8.**
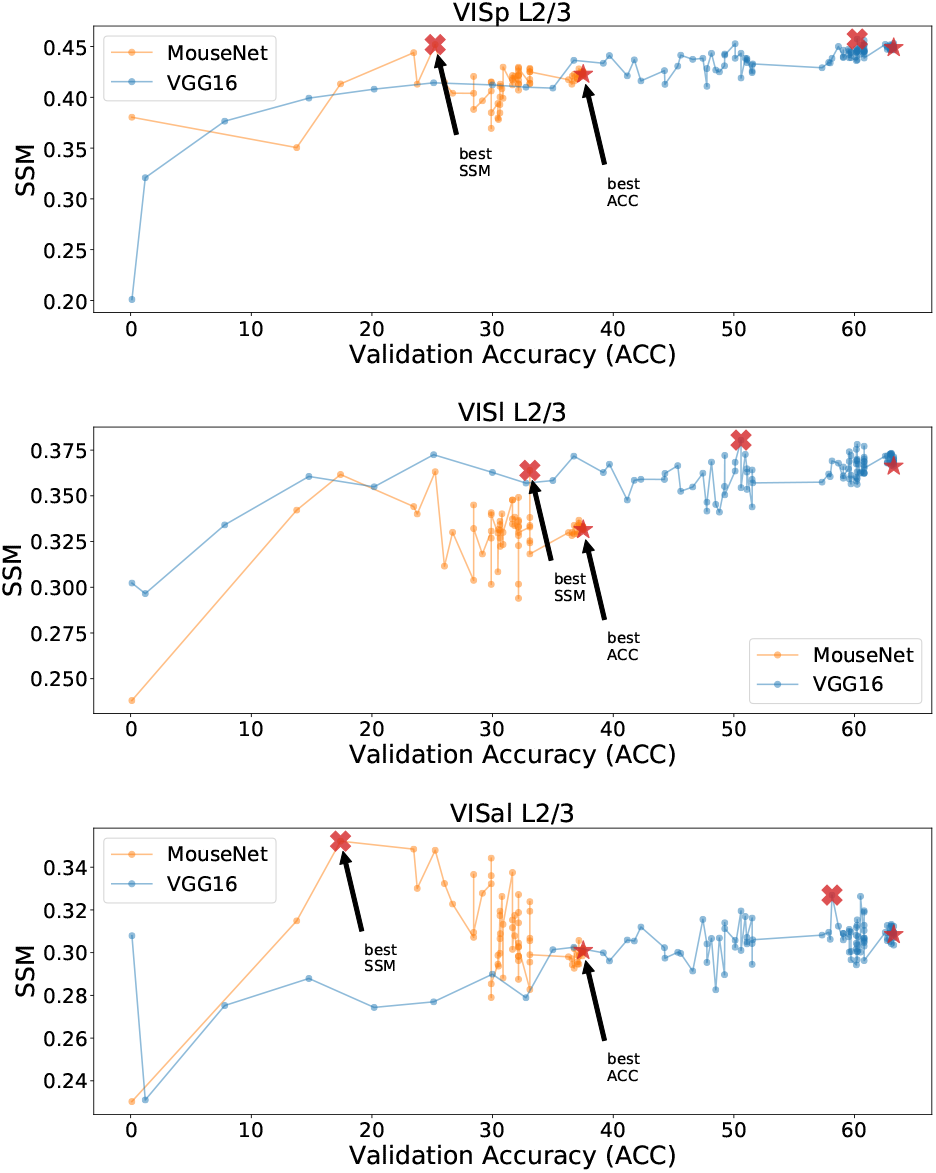
Functional similarity and validation accuracy during the training process. Each row compares models with a different brain area. We show one instance of MouseNet and VGG16 during their training process, where each dot represents the best layer’s SSM of one model at a certain epoch to the specified brain area. The clear jumps of validation accuracy occurred when the learning rate is reduced.

#### Task training with the MouseNet architecture increases the similarity of other functional properties to the mouse brain

As mentioned above, the SSM metric compares functional representations, based on activities of the whole neural population in a given model layer and a set of recordings from a given brain area. For a complementary view of the effect of task training on MouseNet representations, and of the role of its architecture, we can also study the statistical distributions of single neuron functional properties, such as orientation selectivity and lifetime sparseness [26].

Lifetime sparseness measures the selectivity of a neuron’s mean response to different stimulus condition, defined as [26,57]

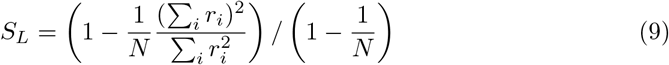

where *N* is the number of stimulus conditions and *r_i_* is the response of the neuron to stimulus condition *i* averaged across trials. A neuron that responds strongly to only a few stimuli will have a lifetime sparseness close to 1, whereas a neuron that responds broadly to many stimuli will have a lower lifetime sparseness. The statistical distribution of lifetime sparseness for the mouse data on natural scene stimuli and for all the units in trained/untrained MouseNet and VGG16 models, responding to the same natural scene stimuli as in the Allen Brain Observatory, are shown in Fig. 9 (top row). This demonstrates clearly that training on the image classification task makes the distribution of lifetime sparseness values much closer to the mouse brain data for MouseNet, but not as much for VGG16.

**Fig 9.**
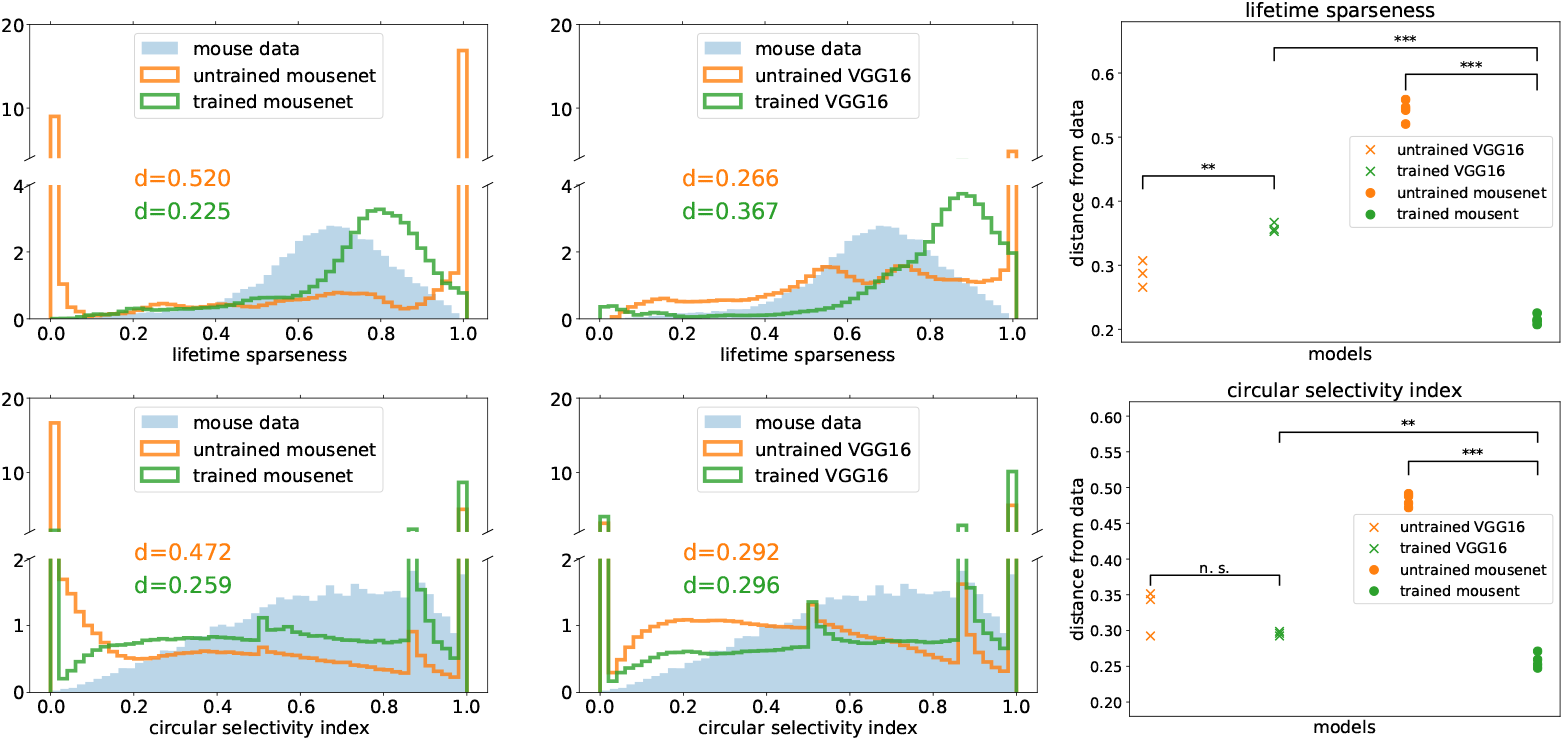
Distributions of lifetime sparseness (top row) and circular selectivity index (bottom row) for all the units in the models and all the neurons in the mouse data. The distributions of all units in one instance of trained/untrained MouseNet (first column) and VGG16 (second column) are plotted along with mouse data, with the Jensen-Shannon distances between the models and the data annotated. The Jensen-Shannon distances between multiple instances of models and the mouse data are summarized in the third column. Black stars denote the statistical significance of two-sample t-test between the mean of the model instances (one star: *p* < 0.05, two stars: *p* < 0.01, three stars: *p* < 0.001).

Similarly, we can study the orientation selectivity of individual neurons by using the static grating stimuli in the Allen Brain Observatory dataset. Specifically, we calculate the circular selectivity index (which is one minus the circular variance defined in [58]), defined as

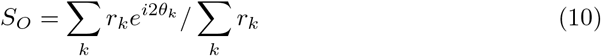

where *r_k_* is the response of the neuron to a grating with angle *θ_k_* averaged across trials. A neuron that responds strongly to only one direction will have circular selectivity index close to 1, whereas a neuron that responds broadly to many directions will have lower circular selectivity index. The statistical distributions of the circular selectivity index, for the mouse data with static grating stimuli and for trained/untrained MouseNet and VGG16 models with the same stimuli, are shown in Fig. 9 (bottom row). As for the case of lifetime sparsity above, task training changes the distribution of selectivity values. These distributions, after training, are closer to the mouse brain data for the MouseNet networks than for the VGG, once again showing how the more specifically matched architecture of MouseNet can lead to more similar model responses to the biological brain. Note that the spikier distributions of the models result from the deterministic nature of the models in contrast to the noisier brain data in response to the (only) six total static grating directions. If we were to simulate neural noise in the model responses, it would smooth the distributions, resulting in closer approximation of the data, as we show in Fig 13 in S1 Fig).

Taken together, these results show how the MouseNet model can be used to explore the impact of task training on a variety of response statistics that are commonly computed in physiology studies, and that those defined on individual neurons can demonstrate complementary and in some cases more dramatic changes with training than those averaged over entire populations.

#### Task training diversifies functional representation among MouseNet layers

Finally, we study how task training and network architecture affect the general ‘geometric’ layout of models’ representations, separately from their similarity to representations in the mouse brain data. To do this, we calculate the SSM values between every pair of layers from both trained/untrained MouseNet and VGG16, and visualize them in two dimensional space via a metric multidimensional scaling algorithm [59,60]. The results are shown in Fig.10. For VGG16, we see that representations in layers are clustered together both before and after training. By contrast, for MouseNet the representations become much more diversified after training. We hypothesize that it is the parallel architecture of MouseNet that leads it to learn this more diversified representation as it solves the image classification task. Further examinations of the various pathways and model instances show that different pathways are learning quite different representations (Fig 14 in S1 Fig), and that these qualitative results are consistent across multiple instances of MouseNet models (Fig 15 in S1 Fig. Unraveling any specific functions of each pathway, in this task or in others, is an opportunity left for future studies.

**Fig 10.**
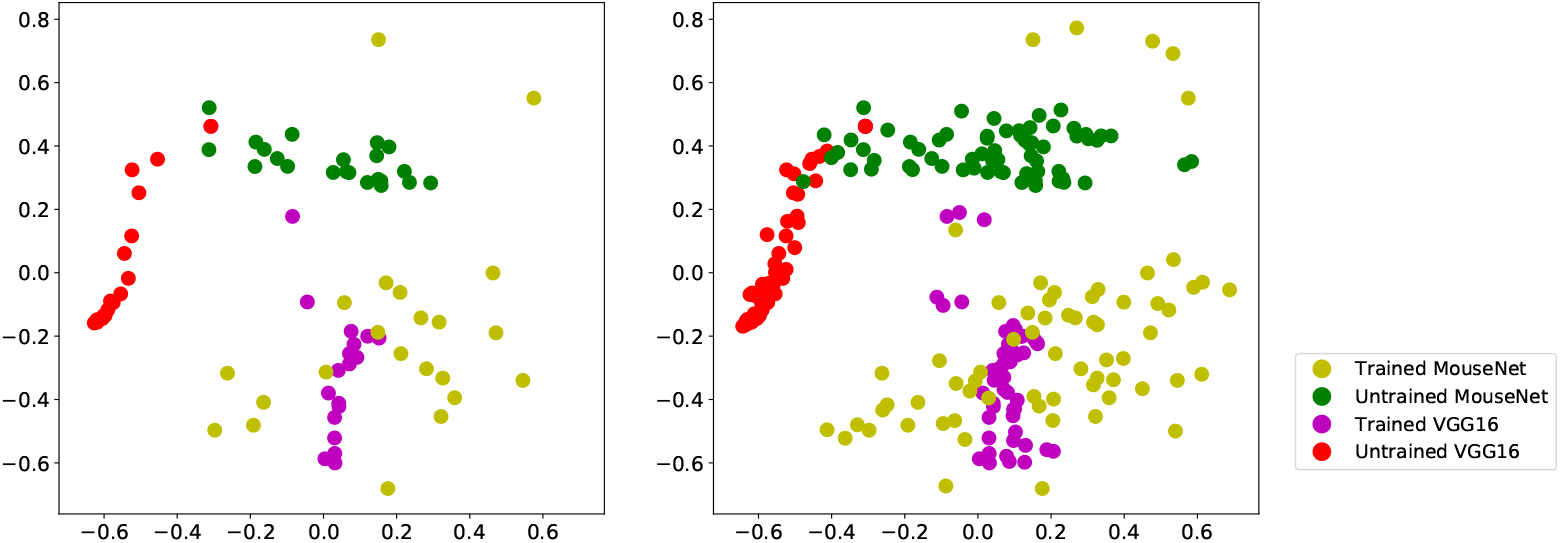
Visualization of all layers from one instance (left) and three instances (right) of trained/untrained MouseNet and VGG16. Each dot represents a layer from a certain model instance. The position of the dots are the two-dimensional projection from the multidimensional scaling algorithm, with the distance measure defined as one minus the SSM value.

## Discussion

Task-optimized deep networks show promise for brain modelling, because they are functionally sophisticated, and they often develop internal representations that overlap strongly with representations in the brain [8–15]. While deep network architectures are originally loosely inspired by the brain, there has been an extensive empirical exploration of the effects of architectural features in machine learning, in directions often independent from neuroscience. In parallel, however, a great deal more has been learned about the architecture of the biological brain, with that of the mouse brain having been been particularly well characterized.

We have developed MouseNet, a deep network architecture that is consistent with a wide range of data on mouse visual cortex, including data from tract-tracing studies and studies of local connection statistics. While standard deep networks have provided useful points of comparison with neurobiological systems, in the long term more biologically realistic deep networks may enable more specific comparisons with the brain, including comparisons between homologous groups of neurons, modeling of specific lesions, and analysis of functional differences between brain areas and pathways.

Using image classification as a working example, we use MouseNet to investigate using the task-training approach to model the functional representations in the mouse brain. An aspect of special interest is whether training on this task drives the representations in the model to be closer to those recorded from the real mouse brain, in comparison to representations in untrained versions of the MouseNet model or in generic deep networks. Using recordings from the large-scale Allen Brain Observatory survey, we find – consistent with the literature [8,9] for other model species and systems – that training on an image classification task does drive MouseNet representations to better resemble those of the real data. However, this increase of functional similarity is not necessarily strictly monotonic with task performance. In our experiments we see the SSM correlation with the Brain Observatory responses saturating or even maximizing well before we achieve maximum accuracy on task performance. This is true for both MouseNet and VGG16.

Within the task-training paradigm, these results suggest that the specific image classification task we used, and perhaps image classification overall, is not the appropriate task for describing what the mouse visual cortex has learned and developed to compute. Nonetheless, MouseNet is an important reference to studies in more established species, which rely on comparisons of the ventral stream with architectures designed for object recognition. Although we know rodents are capable of performing tasks that require visual object discrimination, mouse ethology suggests that alternate computations are more important for the mouse visual system, such as motion tracking, predation, and predator avoidance. A promising future direction is to use task-training of the MouseNet model, together with the metrics tested here, to develop more realistic tasks and stimuli that may lead to more closely matched representations.

In sum this work links anatomical and physiological data to task-driven CNN models, providing a road map for developing better task-driven models of the biological brain. It opens the door to building more detailed structures into the model, such as adding further brain areas as well as adding recurrence and using different inputs and readouts for different pathways. Incorporating new anatomical data is also easy within this framework. By making our code publicly available, and illustrating the model’s success and failures in matching representations using well-studied metrics and tasks, we hope to facilitate future research along these lines.

## Acknowledgments

We thank Tianqi Chen, Blake Richards, Shahab Bakhtiari, Graham Taylor for helpful discussions and suggestions on the manuscript. We thank Saskia de Vries, Hannah Choi, Kameron Decker Harris, Daniel Zdeblick, Timothy Lillicrap, Julie Harris, Severine Durand, and Jun Zhuang for helpful discussions. We thank the Allen Institute for Brain Science founder, Paul G. Allen, for his vision, encouragement, and support. We acknowledge the NIH Graduate training grant in neural computation and engineering (R90DA033461). Research reported in this publication was supported by the National Institute of Neurological Disorders and Stroke and the National Institute of Biomedical Imaging and Bioengineering at the National Institutes of Health under Award Number R01EB026908. The content is solely the responsibility of the authors and does not necessarily represent the official views of the National Institutes of Health.

## Footnotes

1 https://bbp.epfl.ch/nexus/cell-atlas/

2 https://github.com/pytorch/examples/tree/master/imagenet

## Supporting information

S1 Table.

**Table 12.** The estimated 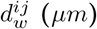 and 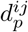 for interareal connections.

| Source | Target | $d_w^{ij}$ ($\mu m$) | $d_p^{ij}$ |
| --- | --- | --- | --- |
| VISp4 | VISal4 | 277.1 | 0.039 |
|  | VISl4 | 313.2 | 0.030 |
|  | VISli4 | 296.7 | 0.032 |
|  | VISpl4 | 290.6 | 0.032 |
|  | VISrl4 | 306.7 | 0.032 |
|  | VISpor4 | 276.8 | 0.013 |
| VISp2/3 | VISal4 | 266.1 | 0.063 |
|  | VISl4 | 325.8 | 0.038 |
|  | VISli4 | 303.3 | 0.045 |
|  | VISpl4 | 284.2 | 0.047 |
|  | VISrl4 | 339.4 | 0.032 |
|  | VISpor4 | 307.4 | 0.013 |
| VISp5 | VISal4 | 239.0 | 0.064 |
|  | VISl4 | 311.5 | 0.042 |
|  | VISli4 | 314.3 | 0.043 |
|  | VISpl4 | 278.3 | 0.053 |
|  | VISrl4 | 311.4 | 0.042 |
|  | VISpor4 | 298.3 | 0.016 |
| VISal4 | VISpor4 | 37.98 | 0.551 |
| VISal2/3 |  | 13.81 | 4.362 |
| VISal5 |  | 14.87 | 4.213 |
| VISl4 |  | 204.7 | 0.014 |
| VISl2/3 |  | 210.6 | 0.016 |
| VISl5 |  | 215.4 | 0.019 |
| VISli4 |  | 155.1 | 0.017 |
| VISli2/3 |  | 169.3 | 0.022 |
| VISli5 |  | 148.4 | 0.028 |
| VISpl4 |  | 22.2 | 0.190 |
| VISpl2/3 |  | 59.5 | 0.054 |
| VISpl5 |  | 54.4 | 0.079 |
| VISrl4 |  | 97.2 | 0.074 |
| VISrl2/3 |  | 105.4 | 0.064 |
| VISrl5 |  | 110.4 | 0.064 |

**Table 13.**
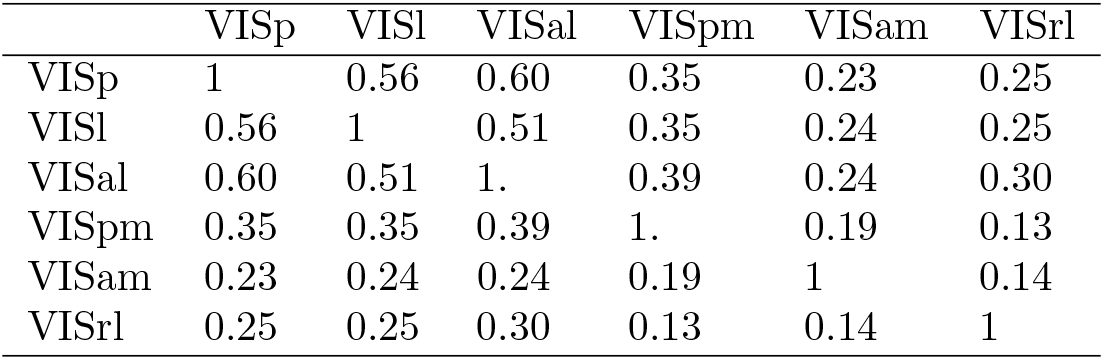
SSM values between mouse visual cortex areas. Note that even with the neural sub-sampling issue [27], the similarity values between VISp, VISl, and VISal are much higher than they are with the CNN models.

|  | VISp | VISl | VISal | VISpm | VISam | VISrl |
| --- | --- | --- | --- | --- | --- | --- |
| VISp | 1 | 0.56 | 0.60 | 0.35 | 0.23 | 0.25 |
| VISl | 0.56 | 1 | 0.51 | 0.35 | 0.24 | 0.25 |
| VISal | 0.60 | 0.51 | 1. | 0.39 | 0.24 | 0.30 |
| VISpm | 0.35 | 0.35 | 0.39 | 1. | 0.19 | 0.13 |
| VISam | 0.23 | 0.24 | 0.24 | 0.19 | 1 | 0.14 |
| VISrl | 0.25 | 0.25 | 0.30 | 0.13 | 0.14 | 1 |

S1 Fig.

**Fig 11.**
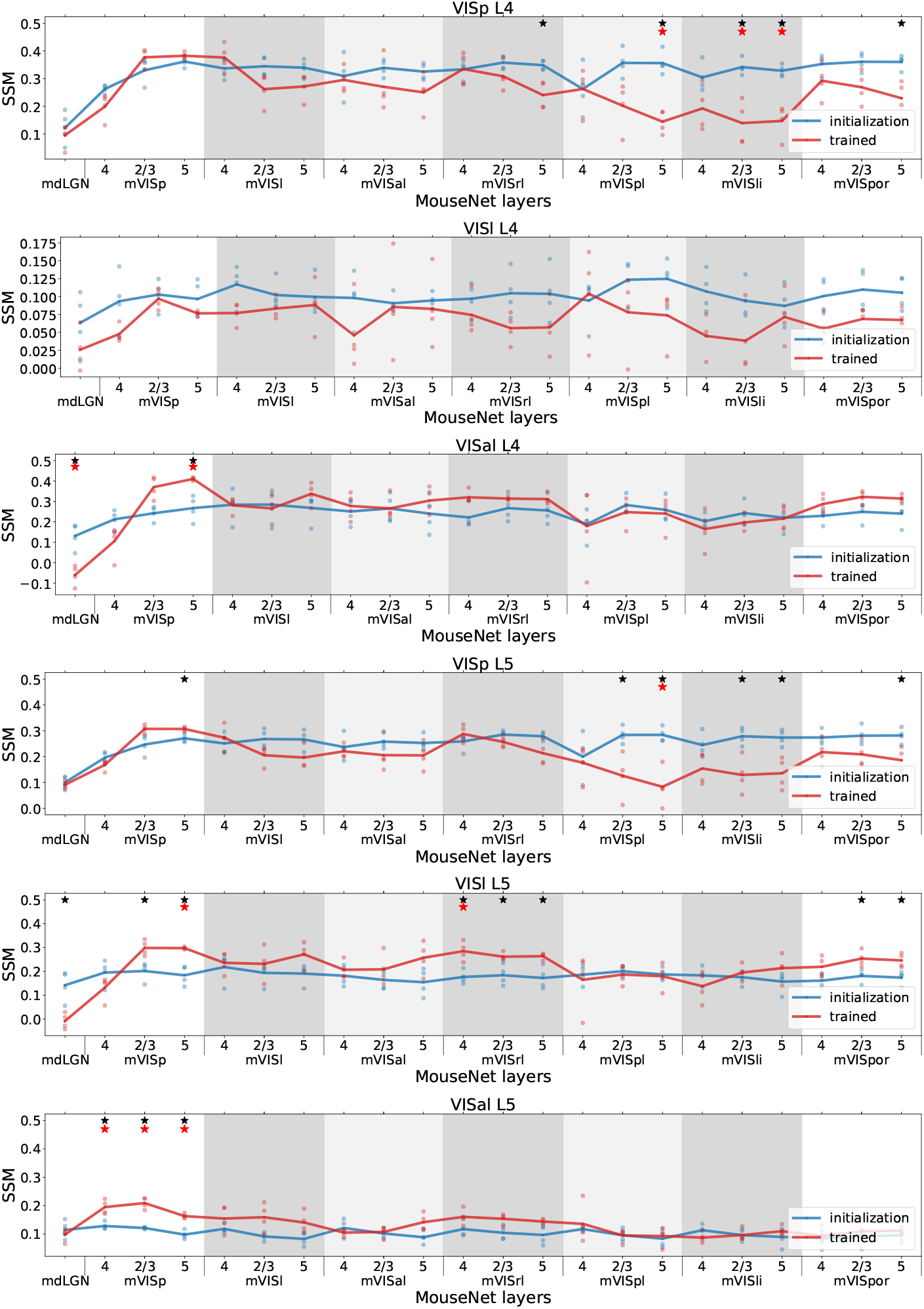
SSM between data in VISp(top)/VISl(middle)/VISal(bottom) L4 and L5 and all layers in the MouseNet before(blue) and after training(red). Each line corresponds to the mean of 4 different MouseNet instances trained from different initialization weights (dots). The x axis includes all the layers in the model in a serial way. The five parallel secondary visual area pathways in the model are in shaded grey background. Black stars denote the the pvalues of two-sample t-test with Benjamini/Hochberg correction of 22 comparisons within one brain area is less than 0.05; Red stars denote the pvalues of two-sample t-test with Benjamini/Hochberg correction of all 9×22 comparisons across all 9 brain areas is less than 0.05.).

**Fig 12.**
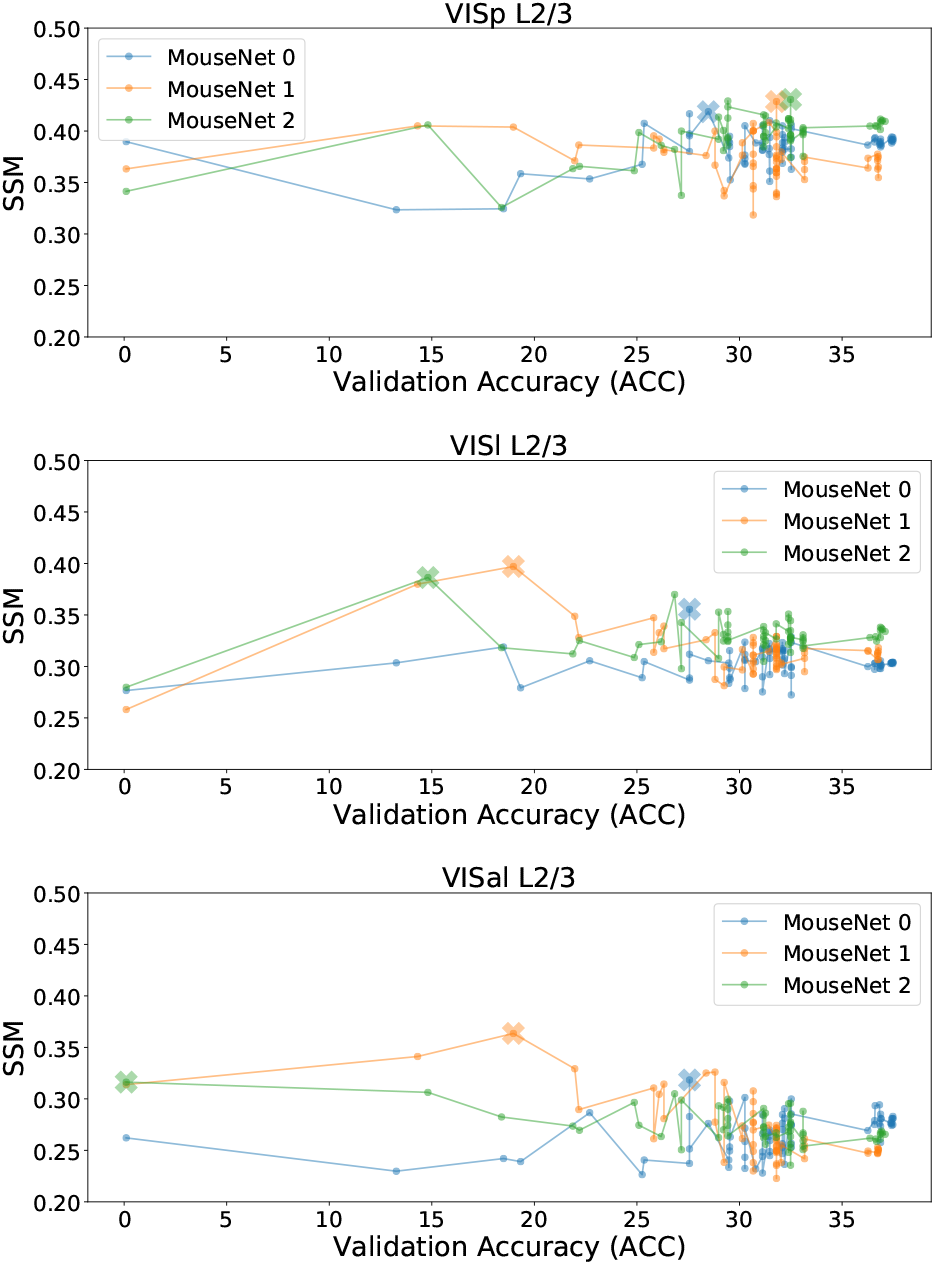
Functional similarity and validation accuracy during the training process for multiple MouseNet instances. Each row compares models with a different brain area. We show three instances of MouseNet during their training process. Each dot represents the best layer’s SSM of one instance at a certain epoch to the specified brain area, with each instance’s highest achieved SSM during training process marked by a cross. The clear jumps of validation accuracy occurred when we reduced the learning rate.

**Fig 13.**
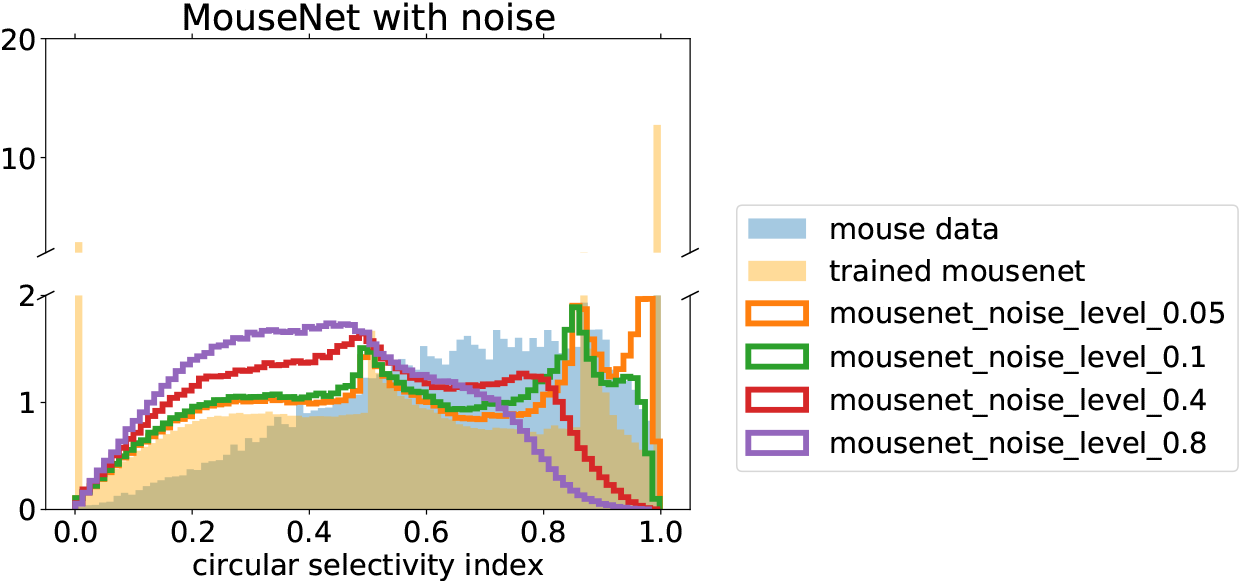
Distribution of circular selectivity index for all the units in trained MouseNet with different levels of noise added. The noise is added to the activations of each layer as a half-normal distribution with a standard deviation of the specified noise level multiplied by the mean activation across all units for that layer. This results shows that circular selectivity index distribution can be smoothed out by adding noise to the deterministic MouseNet model.

**Fig 14.**
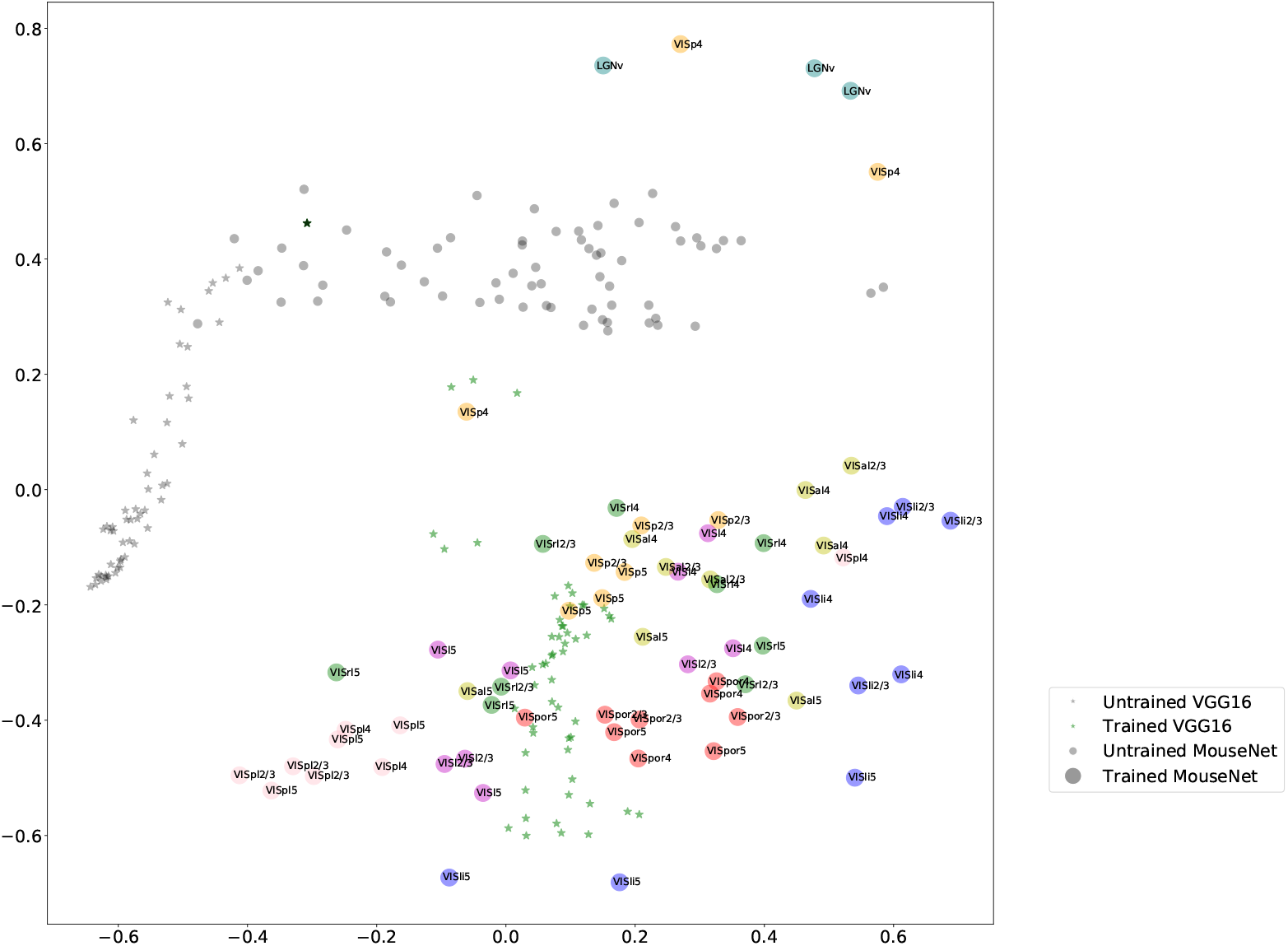
Visualization of all layers of trained/untrained MouseNet and VGG16, for three instances (colored coded by areas). Each dot represents a layer from a certain model instance. The position of the dots are the two-dimensional projection from the multidimensional scaling algorithm, with the distance measure defined as one minus the SSM value. The layers from three instances of trained MouseNet are color coded by their area names, and annotated with their region names. This result shows that different pathways in the MouseNet have learned distinct representations after training.

**Fig 15.**
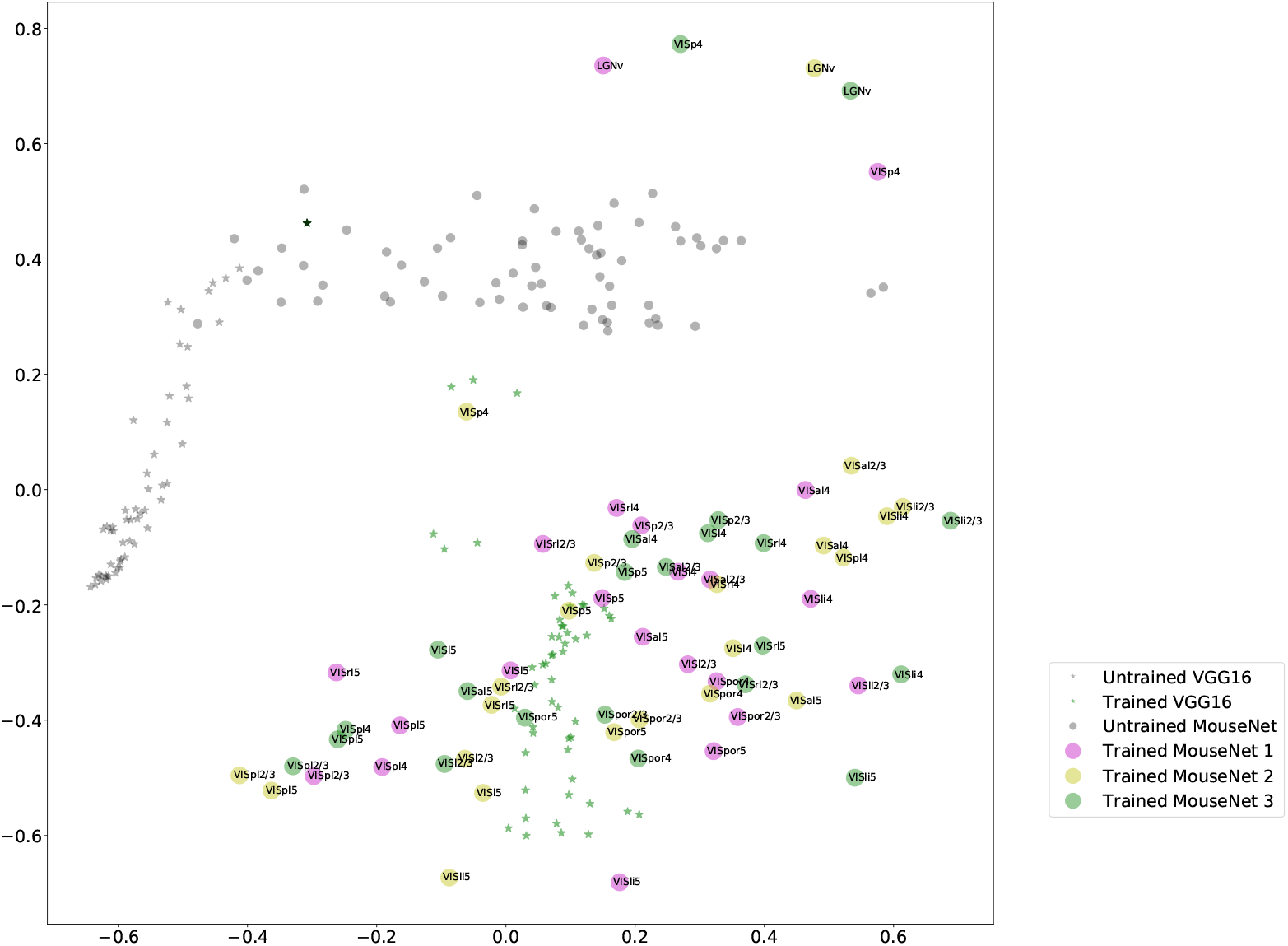
Visualization of all layers of trained/untrained MouseNet and VGG16, for three instances (colored coded by instance). Each dot represents a layer from a certain model instance. The position of the dots are the two-dimensional projection from the multidimensional scaling algorithm, with the distance measure defined as one minus the SSM value. The layers from three instances of trained MouseNet are color coded by their corresponding model instance. This result shows that training diversified the representations of all the three instances of MouseNet starting from different initialization states.

